# KAP1 is an antiparallel dimer with a natively functional asymmetry

**DOI:** 10.1101/553511

**Authors:** Giulia Fonti, Maria J. Marcaida, Louise C. Bryan, Sylvain Traeger, Alexandra S. Kalantzi, Pierre-Yves J.L. Helleboid, Davide Demurtas, Mark D. Tully, Sergei Grudinin, Didier Trono, Beat Fierz, Matteo Dal Peraro

**Affiliations:** Institute of Bioengineering, School of Life Sciences, Ecole Polytechnique Fédérale de Lausanne, Lausanne, 1015 Switzerland; Institute of Chemical Sciences and Engineering, School of Basic Sciences, Ecole Polytechnique Fédérale de Lausanne, Lausanne, 1015 Switzerland; Global Health Institute, School of Life Sciences, Ecole Polytechnique Fédérale de Lausanne, Lausanne, 1015 Switzerland; Interdisciplinary Centre for Electron Microscopy, Ecole Polytechnique Fédérale de Lausanne, Lausanne, 1015 Switzerland; European Synchrotron Radiation Facility (ESRF), 38043 Grenoble, France; University Grenoble Alpes, CNRS, Inria, Grenoble INP, LJK, 38000 Grenoble, France

## Abstract

KAP1 (KRAB-domain associated protein 1) plays a fundamental role in regulating gene expression in mammalian cells by recruiting different transcription factors and altering the chromatin state. In doing so, KAP1 acts both as a platform for macromolecular interactions and as an E3 SUMO ligase. This work sheds light on the overall organization of the full-length protein combining solution scattering diffraction data, integrative modeling and single-molecule experiments. We show that KAP1 is an elongated antiparallel dimer with a native asymmetry at the C-terminal domain. This conformation supports our finding that the RING domain contributes to KAP1 auto-SUMOylation. Importantly, this intrinsic asymmetry has key functional implications for the KAP1 network of interactions, as the heterochromatin protein 1 (HP1) occupies only one of the two putative HP1 binding sites on the KAP1 dimer, resulting in an unexpected stoichiometry, even in the context of chromatin fibers.

## Introduction

KAP1 - KRAB (Krüppel-associated box)-domain associated protein 1-also known as TIF1ß (Transcription Intermediary Factor 1β) or TRIM28 (Tripartite Motif containing protein 28) is a central regulator that controls the fate of the genetic material by recruiting transcription factors and altering the chromatin environment^1,2^. KAP1 is thus essential for early development^3^ and has been linked to fundamental cellular processes such as differentiation^4,5^, gene silencing^6–9^, transcription regulation^10–13^ and DNA damage response^8,14–19^. Moreover, its involvement in control of behavioral stress and tumorigenesis makes it an attractive therapeutic target^20–27^.

KAP1 belongs to the superfamily of the tripartite motif-containing (TRIM) proteins that includes more than 60 members in humans with variable C-terminal domains^28^. The TRIM family is defined by the presence of a highly-conserved N-terminal tripartite motif known as RBCC consisting of a RING (Really Interesting New Gene) finger domain, one or two B-box domains (B_1_ and B_2_) and a long coiled-coil (CC)^28^ (**Figure 1a**). The RING domain contains a regular arrangement of cysteine and histidine residues that coordinate two zinc ions tetrahedrally in a unique “cross-brace” fold^29^ and acts as a E3 SUMO (Small Ubiquitin MOdifier) and E3 Ubiquitin ligase^30–32^. The B-box domain shares the RING domain fold and may bind one or two zinc ions^33^. The CC of KAP1 is estimated to be very long (~200 Å) and together with the B_2_ is likely used to mediate protein-protein interactions^34^.

**Figure 1.**
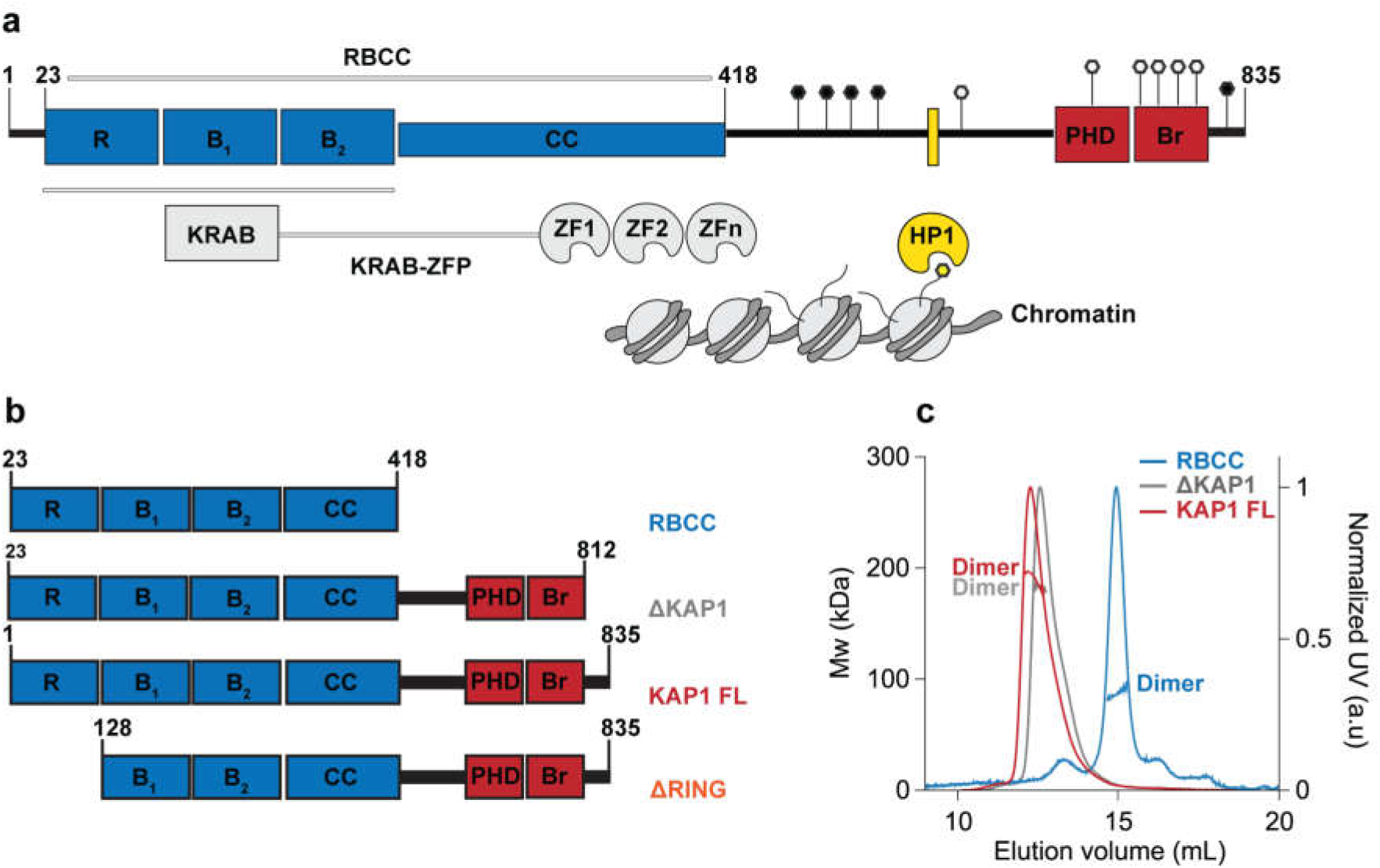
Oligomerization state of KAP1. **a.** KAP1 sequence architecture. The different KAP1 domains are reported on top. The structures of the B-box 2 (residues 201-250) and the PHD-Br domain (residues 624-812) have been solved by X-ray crystallography and NMR, respectively (PDBs 2YVR and 2RO1^40^). Key residues affected by post-translational modifications are also highlighted (phosphorylation sites in black and SUMOylation in white). **b.** Schematic of the different constructs used in this study. **c.** SEC-MALS analyses of the KAP1 constructs show that they are all dimers. The traces are colored according to the KAP1 construct and show the normalized elution profile measured at 280 nm (right axis) and the calculated molecular weight of the selected peaks in kDa (left axis): 88 kDa for the RBCC domain, 183 kDa for ΔKAP1 and 190 kDa for KAP1 FL.

KAP1 is a member of the TRIM C-VI subfamily, together with TRIM24 and TRIM33, characterized by the presence of a tandem plant homeodomain (PHD) and bromodomain (Br) typically involved in the recognition of various histones modifications^35,36^. However, the C-terminal tandem PHD-Br domain of KAP1 shows a unique function acting as an E3 SUMO ligase, promoting both the auto-SUMOylation of the protein^37^ and the SUMOylation of other substrates^38,39^. The NMR structure of the KAP1 PHD-Br domain elucidated how the two domains cooperate as one E3 SUMO ligase unit^40^. The auto-SUMOylation of the C-terminal PHD-Br domain is necessary for the binding of KAP1 to the chromatin remodeling enzymes such as the methyltransferase SET domain bifurcated 1 (SETDB1), the nucleosome remodeling and histone deacetylation complex (NuRD), the histone deacetylase (HDAC), the nuclear co-repressor (N-CoR), inducing the deposition of post-translational modifications (PTMs) and consequently the creation of an heterochromatin environment^41–45^ (**Figure 1a**).

The N-terminal domain and the C-terminal PHD-Br domain are connected by a long loop of ~200 amino acids without any predicted structure, where the heterochromatin protein 1 (HP1) is recruited by KAP1 on a specific binding domain (HP1BD, **Figure 1a**)^46,47^. HP1 is a transcriptional repressor that directly mediates the formation of higher order chromatin structures^48,49^, characterized by the presence of an N-terminal chromo domain (CD) and a C-terminal chromo shadow domain (CSD), linked by an unstructured hinge region. The CD binds directly the methylated lysine 9 of histone H3, while the CSD forms a symmetrical dimer that mediates interactions with other proteins by recognizing a PXVXL penta-peptide motif (where X is any amino acid)^50^. The interaction with HP1 is crucial for KAP1 to regulate the chromatin state^46^, carry out its gene silencing function^51^ and respond to DNA damage^17^.

KAP1 mediates gene silencing of transposable elements by recruiting KRAB-zinc finger proteins (KRAB-ZFPs), which constitute the largest family of TFs with over 350 members and which are present only in tetrapod vertebrates^52,53^. Their function is mainly unknown, but some members have been involved in diverse processes such as embryonic development, tissue-specific gene expression, and cancer progression^54^. KRAB-ZFPs recruit KAP1 to specific genetic loci, via the interaction between the RBCC and the KRAB repression module^9,34,54^, in turn, KAP1 recruits chromatin remodeling enzymes to tri-methylated lysine 9 of histone H3. This modification, along with the DNA compaction caused by HP1 binding, creates a heterochromatin environment silencing gene expression^7^.

Despite its emerging fundamental importance, KAP1 and its interaction with binding partners are still poorly characterized from a structural standpoint. Crystal structures of the CC with or without additional B-box or C-terminal domains have been solved for some TRIM family members (TRIM25^55,56^, TRIM5^57,58^, TRIM20^59^ and TRIM69^60^) showing that the RBCC is an elongated domain with an antiparallel homodimer conformation. However, the dimeric nature of the RBCC in the TRIM family members contradicts published findings suggesting that KAP1 is a homo-trimer in solution and bound to KRAB domains^34^. Here, we present the overall structural organization of human full-length KAP1 in solution by integrating biochemical characterization, small angle X-ray scattering data, molecular modeling and single-molecule experiments. We demonstrate that KAP1 is an elongated antiparallel dimer with a native asymmetry at the C-terminal domains. Based on this conformation, we observe that the N-terminal RING domain is necessary for full auto-SUMOylation. Furthermore, we show that the asymmetric organization is maintained in complex with HP1, which only occupies only one of the two existing binding sites in the KAP1 dimer. We discuss the implications of this asymmetry for HP1 binding kinetics in the chromatin context and in general for KAP1 function.

## Results

### KAP1 is a homodimer in solution

One of the main structural features of the TRIM family members characterized so far is their ability to form homodimers through their coiled-coil (CC) region^55–57,59,61^ (**Figure 1a**). Biochemical and bioinformatics analyses suggest that this organization could be a common feature for all the TRIM family members^56^. Although the homodimer state is the preferred one, other studies have reported the possibility for some TRIM family members to form heterodimers or heterotrimers^62^. Moreover, another layer of complexity is added by the possibility to form high order oligomers mediated by the RING and B-box domains^63^. Unlike the other TRIM family members, KAP1 has been suggested to fold preferentially as a homotrimer alone in solution and in complex with the KRAB domain of KRAB ZFPs^34^. Furthermore, hetero interactions with TRIM24 and further associations as hexamers have been suggested^62^. To shed light on the oligomerization state of KAP1, we performed size exclusion chromatography coupled to multi angle light scattering (SEC-MALS) and sedimentation velocity analytical ultracentrifugation (SV-AUC) experiments. Three different constructs (**Figure 1b**) have been used: (i) the RBCC domain (RBCC, 23-418) mainly responsible for oligomerization, (ii) a KAP1 construct nearly covering the whole sequence (ΔKAP1, 23-812) and (iii) the complete KAP1 full length protein (KAP1 FL, 1-835). SEC-MALS experiments were performed across a concentration range of 10 to 40 μM (**Figure 1c**). No concentration dependence in either elution volume or mass estimation was observed. The resulting molecular weights (Mw_s_) were determined to be 88 kDa for the RBCC domain, 183 kDa for ΔKAP1 and 190 kDa for KAP1 FL, in agreement with dimeric species of expected Mw_s_ of 92 kDa (RBCC), 175 kDa (ΔKAP1) and 182 kDa (KAP1 FL) (**Figure 1c**). Under the concentration range tested, both SEC-MALS and SV-AUC (**Supplementary Figure 1**) did not show the higher order oligomers observed for other TRIM family members, such as TRIM32^61^ and consistently indicated a dimeric conformation for KAP1.

### KAP1 is an antiparallel elongated dimer determined by the coiled-coil domain

In order to gain more information on the structural organization of KAP1 in solution, we performed small angle X-ray scattering (SAXS). Size-exclusion chromatography was coupled in line with SAXS to avoid self-association and aggregation effects during the experiments (**Figure 2a**). While SAXS data were recorded for the three KAP1 constructs, ΔKAP1 and KAP1 FL showed identical results and are reported in **Supplementary Figure 2**. The data were collected across a concentration range of 9 to 15 mg/ml (100-200 μM). The Guinier analysis of the scattering curves showed good linearity indicating neither aggregation nor polydispersity effects and gave an estimated radius of gyration (R_g_) of 83 Å for RBCC, 90 Å for KAP1 FL and 89 Å for ΔKAP1 (**Supplementary Table 1**). Moreover, the value for the cross section R_g_ (R_gc_, **Supplementary Figure 3**) was similar for KAP1 FL (35.8 Å) and ΔKAP1 (38.8 Å), while it was smaller for the construct containing only the N-terminal RBCC domain (20.2 Å) (**Supplementary Table 1**) pointing to an elongated structure for KAP1. Additionally, the SV-AUC experiment on the RBCC construct showed the protein to sediment as a single species with an S_20_,w of 2.3 and a frictional ratio (f/f_o_, where f_o_ is the frictional coefficient of a smooth compact sphere) well above 1.3, indicating the elongated nature of the molecule and confirming the SAXS observation (**Supplementary Figure 1**). Furthermore, the analysis of the Kratky plot, which can detect features typical of multi domain proteins with flexible linkers, indicates a large flexibility of KAP1 (**Supplementary Figure 4**). This observation agrees with the prediction of an unstructured 200 residue-long loop connecting the N-terminal RBCC and the C-terminal PHD-Br domains.

**Figure 2.**
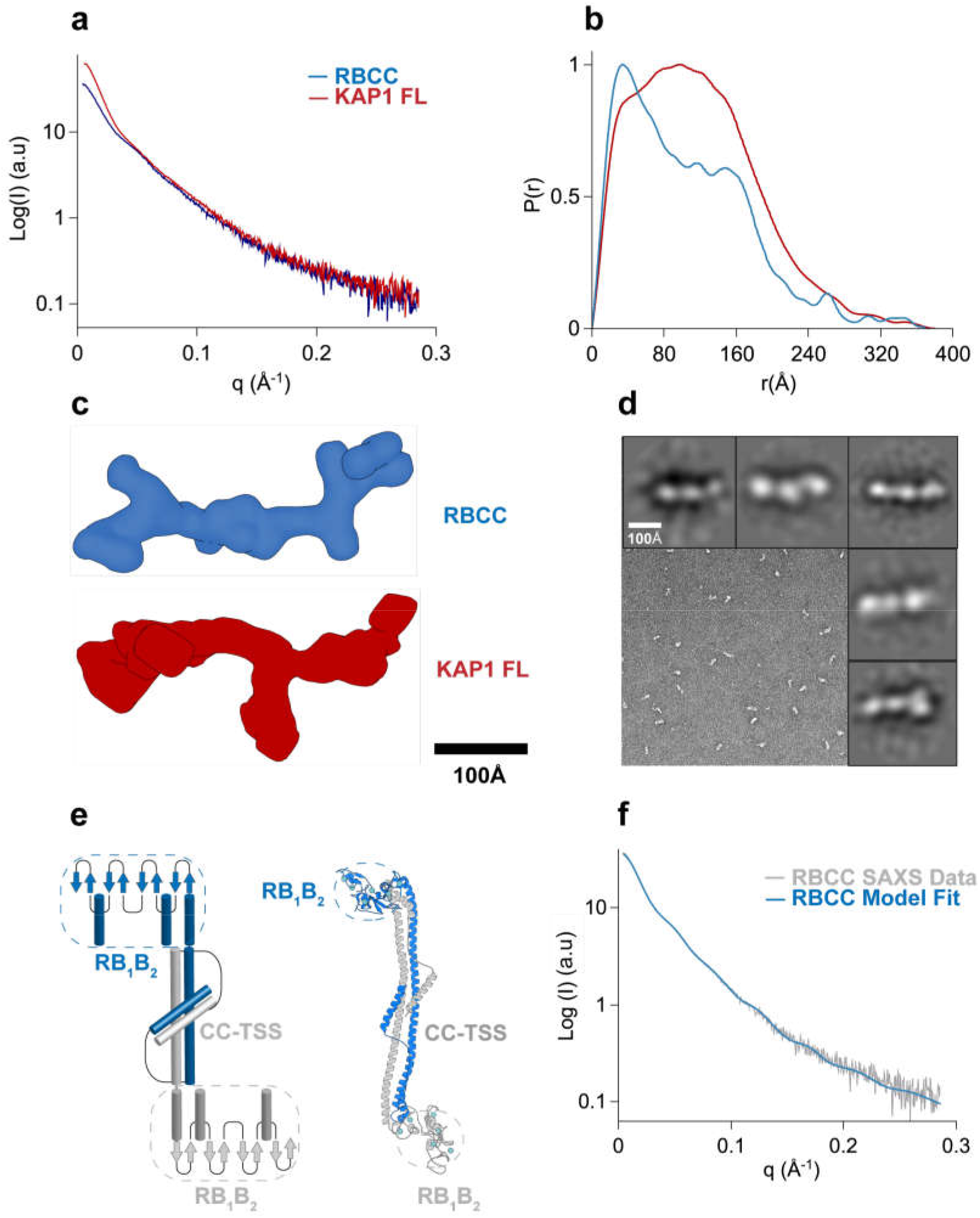
Overall architecture of KAP1. **a.** SAXS scattering curves. **b.** Pair-distance distribution functions. **c.** *Ab initio* bead models created using GASBOR^64^ shown in surface representation. **d.** Representative TEM image of negatively stained KAP1 FL and gallery of 2D class averages. **e.** Schematic representation and atomistic model of the antiparallel RBCC dimer (different protomers are colored in blue and grey). The end of the RBCC contains the TIF1 signature sequence (TSS) domain, which is rich in phenylalanine and tryptophan residues^2^ and is modeled as a helix that packs against the main long CC domain forming overall a 4-helix bundle. **f.** Fit of the calculated scattering profile of the molecular model in (**e**) (in blue) to the SAXS scattering data (in grey) using Pepsi-SAXS^65^ (χ^2^=0.9).

MALS and SAXS analyses reveal that KAP1 is an elongated flexible dimer. Therefore, to gain insight into its mass distribution, we compared the pair distribution functions, *P*(*r*) of the RBCC domain and KAP1 FL. Both show the signature of an elongated molecule with very large maximum dimension (*D_max_*) values and a main peak at a shorter radius (**Figure 2b**). The *P*(*r*) function for the RBCC is clearly bimodal, containing two peaks one centered at 40 Å and the other one at 150 Å with a *D_max_* of 370 Å (**Figure 2b** and **Supplementary Table 1**) suggesting a dumbbell-shaped macromolecule with the RBCC domain arranged in an antiparallel fashion separating the two N-terminal RING/B-box 1/B-box 2 (RB_1_B_2_) modules by its length (160 Å). The *P*(*r*) function of KAP1 FL is also characteristic of an elongated molecule but surprisingly it shows a similar *D_max_* (380 Å) as for the RBCC domain alone. This observation, together with the increase in *R_gc_*, leads to the conclusion that the extra domains present in the full-length protein are not fully extended but are interacting closely with the N-terminal RBCC domain. This conclusion is also supported by the loss of the bimodality in the *P*(*r*) function compatible with a more compact protein in which the extra C-terminal domains interact with the RBCC domains (**Figure 2b**). In addition, *ab initio* bead models calculated directly from the SAXS scattering curves (**Figure 2c**) provided conformations of similar length (~320 Å), but differed width, in agreement with the larger R_gc_ of KAP1 FL.

Transmission electron microscopy was used to visualize negatively stained KAP1 FL to independently measure its size and shape (**Figure 2d**). Rod-like particles could be readily observed in the micrograph when sorted into 2D class averages. In agreement with the SAXS data, the particles are around ~280 Å and globular shapes can be observed along the length of the molecules, which can be associated to the different domains.

The RBCC domain being a dimer with an elongated dumbbell-shaped conformation is only compatible with an antiparallel conformation in which the two tandem RB_1_B_2_ modules are separated by the CC domain (**Figure 2e**). Therefore, we could build an atomistic model of the RBCC domain using homologous TRIM structures as templates. Despite the low sequence homology among family, model building was facilitated by conservation of the canonical left-handed coiled-coil motif and the zinc-coordinating His/Cys pairs of the B-box 2 domain among 54 human TRIM proteins^56^. The RBCC model, fitted to the SAXS data using a χ^2^–minimizing optimization^66^ (Methods and **Figure 2f**) showed an elongated structure of ~310 Å in length, with the RB_1_B_2_ modules separated by ~160 Å, the dimension of the dimeric antiparallel CC domain. Interestingly, when compared to published SAXS data of other TRIM family members, the R_gc_ of the RBCC KAP1 is ~20 Å, remarkably smaller compared to the R_gc_ of RBCC-TRIM25 (~31 Å) and RBCC-TRIM32 (~32 Å)^61^. In the first case, the difference in R_gc_ value can be due to the fact that the RB_1_B_2_ modules of TRIM25 are thought to fold back on the structure of the CC, while for KAP1 can be fitted only at the extremes of the CC. In the second case, RBCC-TRIM32 behaves as a tetramer in solution^61^ while we found KAP1 exclusively as a dimer in solution at the sampled concentrations (**Figure 1c**).

### KAP1 dimers are natively asymmetric in solution

The PHD-Br domain of KAP1 acts as an E3 SUMO ligase^37^, which renders KAP1 unique within the TRIM family. The overall organization of the full length KAP1 is unknown as well as the possible interactions, functional cooperation or structural stabilization between the RBCC and C-terminal domain. Even though *ab initio* reconstruction is well suited for relatively rigid molecules^68,69^, for flexible systems with long flexible linkers it can be very ambiguous. Therefore, to gain deeper insight into the domain organization of KAP1 we devised an integrative modeling strategy^70–72^ that uses nonlinear Cartesian Normal Mode Analysis (NOLB NMA)^66^ to optimally fit the SAXS data.

Briefly, 1000 randomized atomistic models of KAP1 FL were generated based on the dimeric RBCC model (**Figure 2e**) and the structure of the C-terminal PHD-Br domains, where ~200 amino acids between the RBCC and the PHD-Br domains were modeled as a random coil structure (Methods). The NMA technique was used to efficiently explore the configurations of the flexible linkers and rigid domains in a reduced conformational space of dimensionality 60^65^. The final models converged to a structure well consistent with SAXS data (χ^2^ = 1.1±0.3), reducing their R_g_ values from ~94±7 to ~88±2 Å, a value similar to the experimental R_g_ value for KAP1 FL (**Figure 3a** and **Supplementary Table 1**). Relevant distances of the mutual positions of the domains separated by the flexible loops (*dA_i_* and *dB_i_* in **Figure 3b**) were used to characterize the global architecture of the KAP1 FL. As expected, the initial models had a very heterogeneous ensemble of conformations, however after flexible fitting two distinct clusters could be identified (**Figure 3c**). Due to the dimeric nature of KAP1, the two clusters are symmetric and their centroids represent one unique conformation that optimally fits the SAXS data (**Figure 3d,e**). This KAP1 model is characterized by a large displacement (~120 Å) of one PHD-Br domain from the RBiB_2_ region and a small displacement (~60 Å) of the other PHD-Br domain from the remaining RB_1_B_2_. The two PHD-Br domains are never found simultaneously close to the RBCC domains, contrary to what was proposed for the TRIM25 PRYSPRY domain and the NHL repeats of TRIM32^61^. If such conformation is imposed, this induced large distortions of the CC domain and poor data fit (χ^2^ >80). Similarly, flexible fitting never selected conformations where the two C-termini are fully extending away from the RBCC domains (χ^2^>75). Consistently, *P*(*r*) functions calculated a posteriori from models optimally compare to experimental *P*(*r*) only for the centroid model structures (**Supplementary Figure 5**).

**Figure 3.**
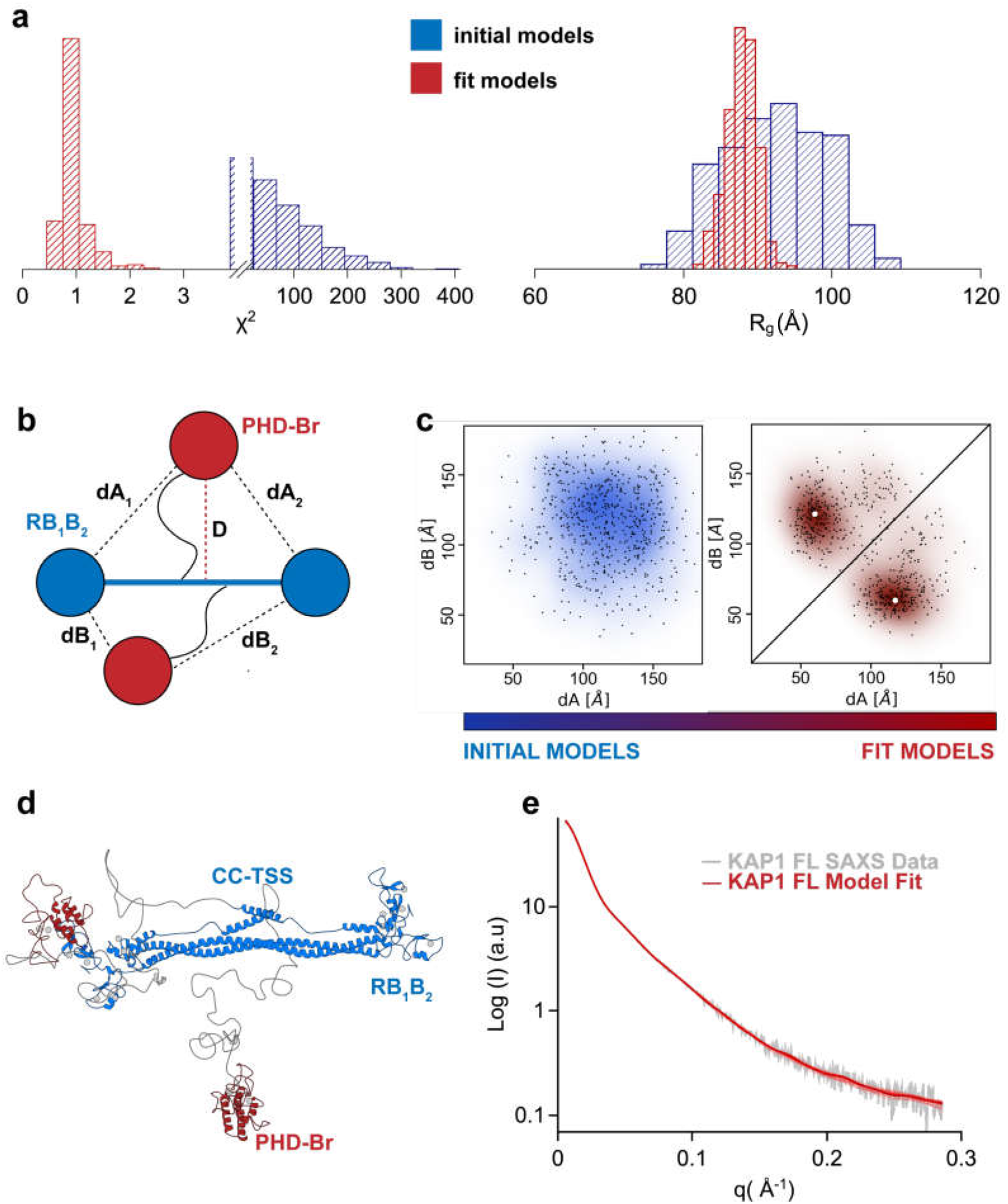
Native asymmetry of KAP1. **a.** χ^2^ (left) and *R_g_* (Å) plots (right) of the initial (blue) vs. final (red) model structures, with final distributions centered around χ^2^=1 and the experimental *R_g_* (90 Å). **b.** Schematic representation of the KAP1 FL dimer where the distances, *D* (maximum initial distance), *dA_1,2_* and *dB_1,2_* are displayed. c. Plot of the initial vs. final models according to their *dA* vs. *dB* distance values. The centroids of the clusters are highlighted by white dots. **d.** Cartoon representation of the centroid model of KAP1 FL with the domain arrangement that best fit the SAXS data. **e.** Comparison between the SAXS scattering profile (grey) and the calculated scattering profile (red) from the centroid structure shown in (**d**) using Pepsi-SAXS (χ =1). The calculated scattering profile for representative models spanning the χ range of [0.8,2] in (**a**) are plotted in red shade on top of the centroid fit.

Therefore, SAXS data strongly indicate that the organization of KAP1 FL is a natively asymmetric dimer, arranged with one of the C-terminal PHD-Br domains closely interacting with one RB_1_B_2_ domain. This conformation has the effect to inhibit the second PHD-Br domain from adopting a similar conformation at the same time. Instead, it remains detached from the elongated core of KAP1, resulting in being more exposed to the bulk and not directly engaging interactions with RB_1_B_2_. We cannot exclude a dynamic equilibrium between two identical conformations in which a PHD-Br domain can swap location affecting the position of the other (i.e. symmetric cluster in **Figure 3c**). Moreover, whether the PHD-Br domain is establishing interactions with the RB_1_B_2_ intra- or inter-molecularly remains an open question.

### KAP1 architecture allows RING-dependent auto-SUMOylation

We asked next if this novel and unexpected conformation of the KAP1 C-terminal domain in solution may have direct implications for its function. The residue C651, located in the PH domain, has been shown to be the key residue for the auto-SUMOylation of the C-terminus^37^ (**Figure 1a**). This post-translational modification is fundamental because it allows KAP1 to interact with chromatin remodeling enzymes inducing the formation of heterochromatin^42,43^. However, previous studies reported that the intact PH domain was necessary for the auto-SUMOylation of many but not all the sites in KAP1^37^, implying the existence of a second catalytic site. Similarly, the RING domain has been shown to strongly interact with the Ubc9 E2 SUMO ligase and be fundamental for the SUMOylation process of KAP1 substrates such as the Interferon Regulatory Factor 7 (IRF7)^31^ and the neurodegenerative disease driving proteins tau and a-synuclein^32^. Therefore, the proximity of one RING to a PHD-Br domain proposed by our SAXS based model (**Figure 3d**) hints to a role for the RING domain in the auto-SUMOylation of KAP1. In order to prove this hypothesis, we performed a SUMOylation assay *in vitro* using purified proteins, where KAP1 acts as an E3 SUMO ligase, auto-SUMOylating itself. We compared the auto-SUMOylation of KAP1 FL, RBCC, KAP1 FL C651A mutant and a deletion mutant missing the N-terminal RING domain (ΔRING) (**Figure 1b**), after having checked that they were properly folded (**Supplementary Figure 6**). As expected, the RBCC domain was not SUMOylated, but differences in the time-dependent SUMOylation of the other variants were observed (**Figure 4**). The C651A mutant showed residual SUMOylation activity with respect to the wild-type protein (40% of FL activity at 120 min, **Supplementary Figure 7**) as previously reported^37^. More importantly, we found that also the ΔRING construct presented a much lower auto-SUMOylation activity than KAP1 FL (~60% at 120 min, **Figure 4** and **Supplementary Figure 7**). Taken together, these results show that not only the PHD but also the N-terminal RING domain is implicated in the auto-SUMOylation process of the C-terminal domain. The conformation discovered by our SAXS-based modeling (**Figure 3d**) is thus consistent with a direct communication between the two terminal domains, which both cooperate in the E3 SUMO ligase activity of KAP1. The RING domain can thus act as a second active site, as it is able to SUMOylate other substrates^31,32^.

**Figure 4.**
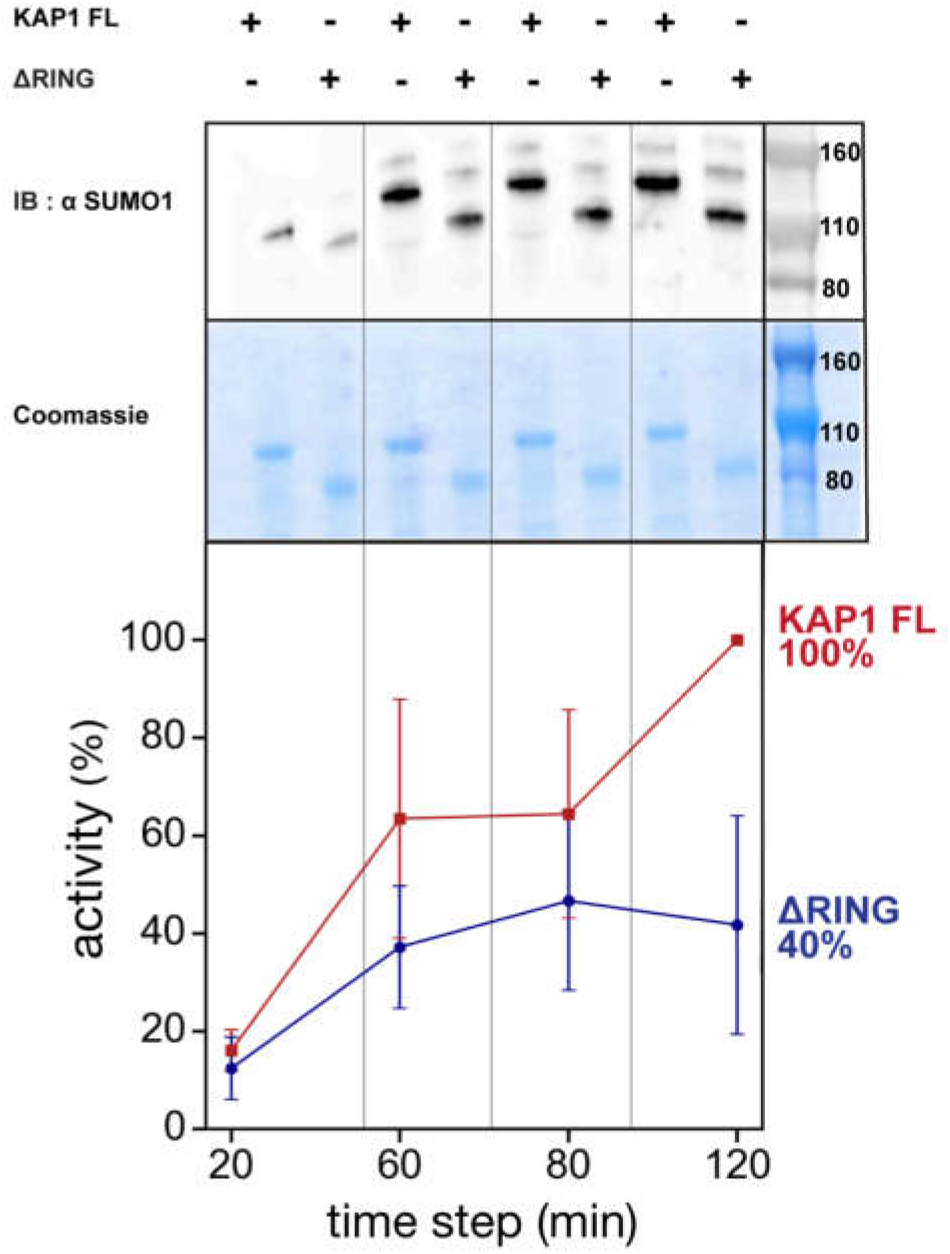
The RING domain contributes to auto-SUMOylation of KAP1. *In vitro* SUMOylation of KAP1, where reactions containing E1, E2 (Ubc9), SUMO1 and KAP1 variants (FL and ΔRING) were incubated for the time indicated (20 to 120 min). The samples were analyzed by western blot with anti-SUMO1 antibody (top) and Coomassie blue staining (center). The intensity of the bands was quantified using ImageJ^75^ and plotted as percentage values of the maximum activity of KAP1 FL (100% at 120 min, bottom). The mean value and standard deviation for each time step are reported for the four technical replicas shown in **Supplementary Figure 7.** Auto-SUMOylation for RBCC, C651A mutant, and controls are also reported in **Supplementary Figure 7.**

Interestingly, RING-mediated auto-SUMOylation has been observed also in another TRIM family member, the protein PML (Promyelocytic leukemia, TRIM19), which is involved in the formation of nuclear structures called PML-nuclear bodies implicated in a variety of cellular processes^73,74^.

### KAP1 asymmetry is functional for recruiting binding partners

To understand if KAP1 structural asymmetry has functional implications when interacting with other binding partners, we explored the direct interaction between KAP1 FL and full length HP1α (HP1α FL). Previous studies have identified a fragment of 15 residues within the central unstructured region of KAP1 as responsible for HP1 binding (residues 483-497, HP1BD, **Figure 1a**)^41,46^. Moreover, mutations of the HP1BD abolish the HP1 binding and significantly reduce the repression activity of KAP1^46^. This interaction, which is independent from post-translational modifications, is mediated by the CSD of HP1 that binds as a dimer to the HP1BD. All previous studies explored the binding between the two proteins using purified fragments, specifically the CSD of HP1 and the HP1BD peptide of KAP1. Here, using full length HP1 and KAP1 we determined their binding affinity by isothermal titration calorimetry (ITC, **Figure 5a**), observing an equilibrium dissociation constant (K_d_) of 162 nM for the KAP1-HP1 complex, value which is in line with previous measurements obtained using protein fragments^46^ and which confirms the strong affinity between the two proteins.

**Figure 5.**
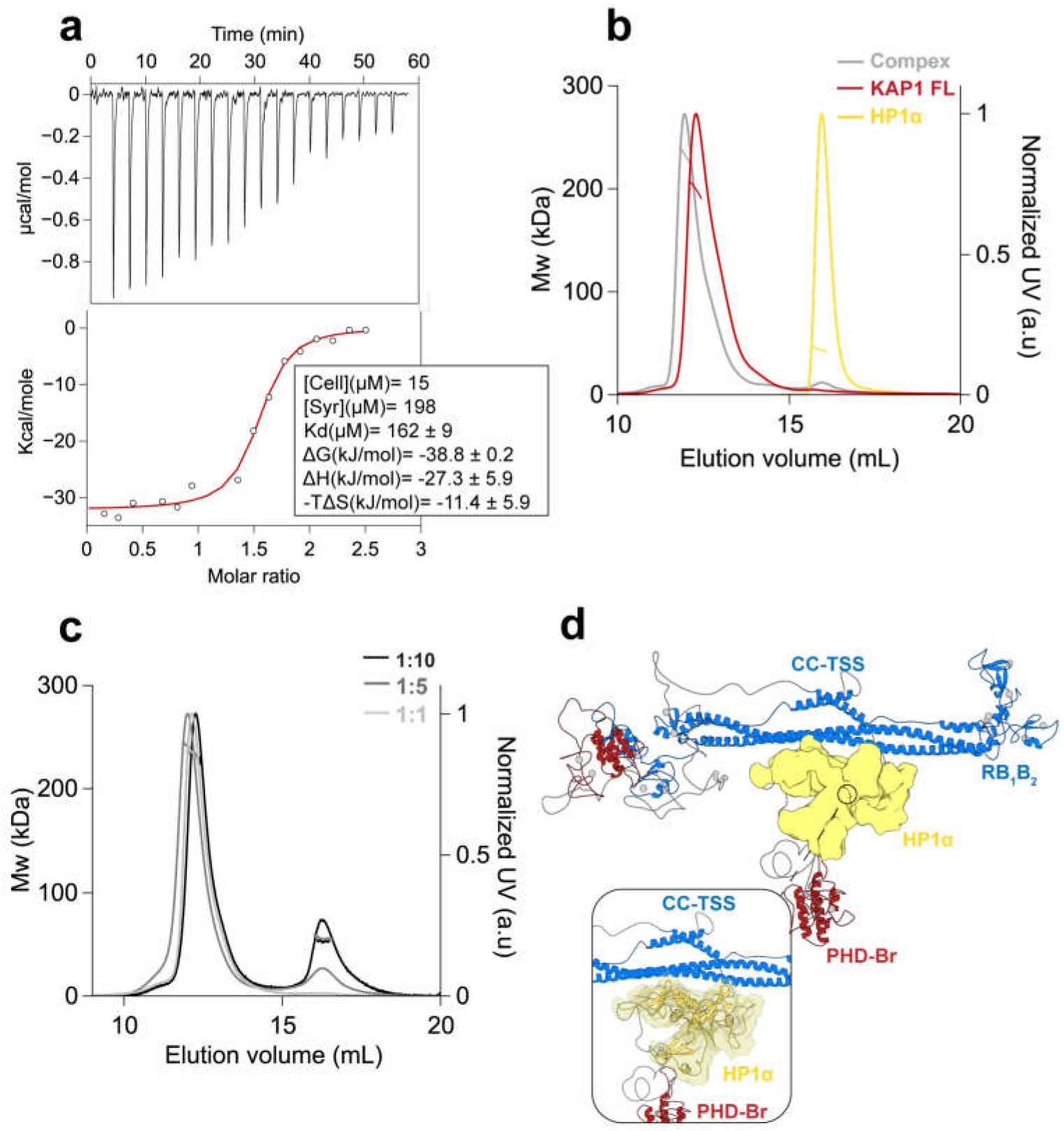
KAP1 dimer binds only one HP1 dimer. **a.** ITC experiment titrating HP1α FL into KAP1 FL measures a tight interaction with a K_d_ of 162 nM. **b.** SEC-MALS analysis comparison between KAP1 FL (red), HP1α FL (yellow), and complex (grey). **c.** SEC-MALS analysis of the complex at different concentration ratios. The estimated masses for the complexes eluting at 11.8 ml when mixtures contained KAP1:HP1 ratios of 1:1, 1:5 and 1:10, were 233 kDa, 239 kDa, and 232 kDa, respectively. The second peak eluting at 16 ml is the excess HP1 dimer with a mass of 45 kDa. **d.** Representative model of the interaction between a KAP1 FL dimer (blue and red) and an HP1 FL dimer (yellow). The CSD of HP1 has been placed as to bind the accessible HP1BD (insert). Further details about the modeling of the HP1 FL dimer are reported in **Supplementary Figure 8.**

Furthermore, we determined the molecular weight of the complex by SEC-MALS, exploring a range of concentrations from 10 μM to 127 μM for KAP1 FL and from 50 μM to 500 μM for HP1α FL, i.e. covering complex stoichiometries ranging from 1:1 to 1:10 in order to saturate binding. The proteins elute as single symmetric peaks with retention volumes of 12 ml for KAP1 FL, 16 ml for HP1α FL and 11.8 ml for the HP1-KAP1 complex (**Figure 5b**), and no concentration dependence in either elution volume or mass estimation was observed (**Figure 5c**). Thus, our measurements estimate a molecular weight of ~231 kDa for the complex, only compatible with a dimer of KAP1 FL (182 kDa) bound to one dimer of HP1α FL (45 kDa), for a 2:2 stoichiometry of the KAP1-HP1 a complex. The expected stoichiometry 2:4, having two HP1α dimers bound to the two HP1BDs of the KAP1 dimer (Mw of ~276 kDa) was never observed. Based on this finding, we speculate that the reason for such unexpected coupling with HP1 has to be related with the asymmetric nature of KAP1, which in turn affects the ability to expose the HP1BD and recruit efficiently HP1 molecules. The asymmetric KAP1 structure has in fact only one HP1BD fully exposed for HP1 interaction, while the second is mostly interacting with the RBCC domain (**Figures 2c, 3d** and **5d**).

The complexity of KAP1-HP1 mediated repression is further enhanced by the flexible conformations of multi-domain proteins involved in protein-protein and protein-chromatin interactions, all timely modulated by post-translational modifications. The study of such systems benefits from the combination of structural information and measurements of the interaction dynamics between components at the single-molecule (sm) level. Total internal reflection fluorescence (smTIRF) imaging approaches can directly observe the chromatin interaction kinetics of individual chromatin binders^76,77^. We used this technique to determine how KAP1 alters the interaction dynamics of HP1α with chromatin fibers, trimethylated at lysine 9 on histone H3 (H3K9me3) to observe the behavior of the complex in the context of chromatin (**Figure 6a**). Here, H3K9me3 containing chromatin fibers (labeled with a fluorescent dye, Atto647N) are immobilized in a flow cell, and fluorescently labeled HP1α (carrying an Atto 532 dye) is injected, in the presence or absence of KAP1. Transient HP1α binding interactions on individual chromatin fibers are then directly observed using smTIRF (**Figure 6b,c**). Compared to the short dwell times of individual HP1α molecules on chromatin (**Figure 6b**), the presence of 100 nM KAP1 resulted in a higher frequency of long (> 4 s) binding events (**Figure 6c**). Analyzing dwell-time histograms (**Figure 6d**) revealed bi-exponential binding kinetics for HP1α (characterized by a fast and slow residence time *τ*_off,1_ and *τ*_off,2_, **Table 1**). It was previously shown that the slow exponential phase is attributed to bivalent chromatin binding by HP1α dimers^76^. KAP1 addition indeed stabilized HP1α chromatin binding, exhibiting a larger percentage of bound molecules at times >1s (**Figure 6d**). Conversely, binding events were less frequent in the presence of KAP1, as indicated in the slower binding kinetics (**Figure 6e**).

**Figure 6.**
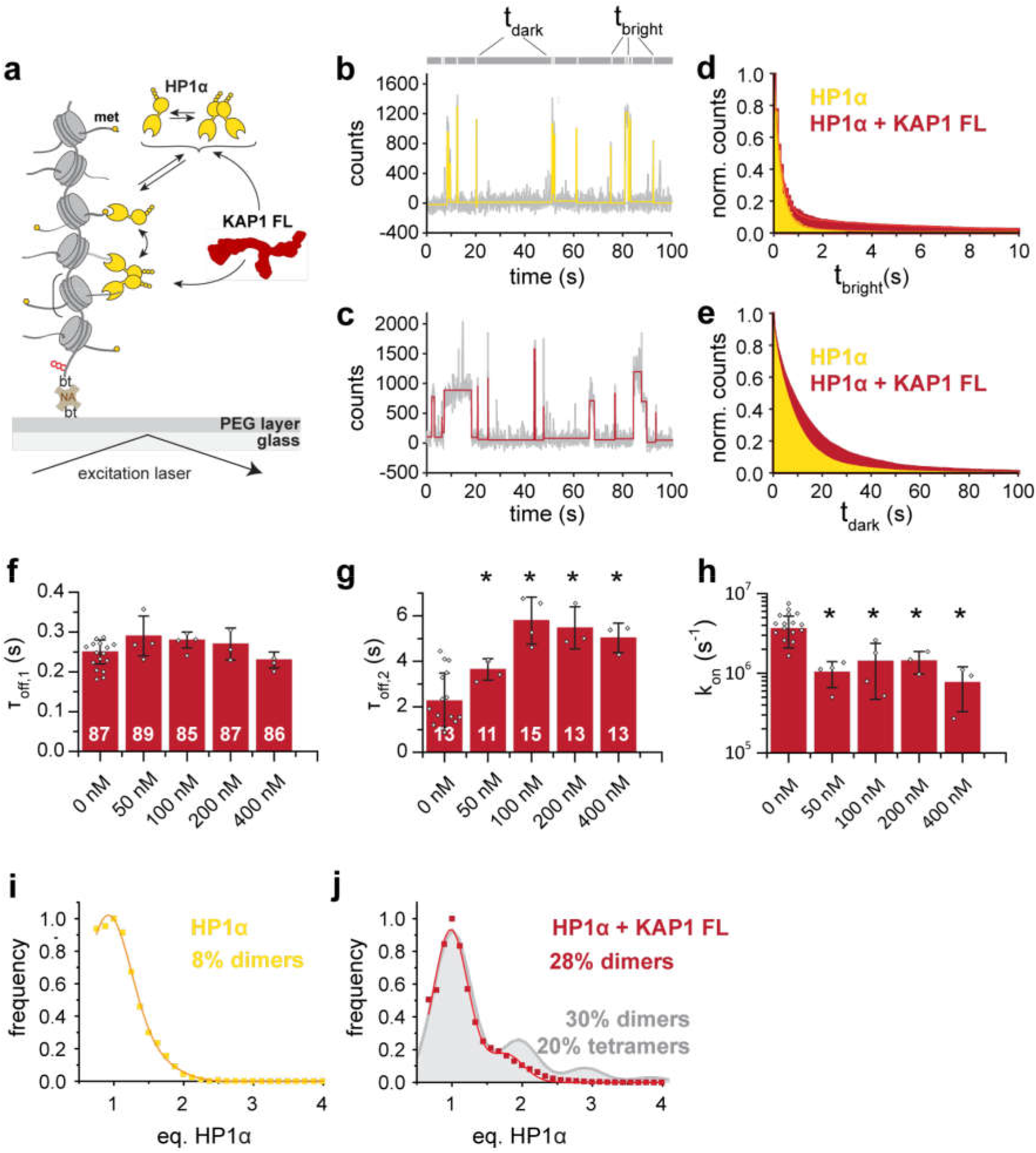
KAP1 stabilizes HP1α-chromatin interactions. **a.** Schematic representation of the smTIRF imaging experimental setup, showing HP1α interacting with chromatin fibers in the presence of KAP1. bt: biotin, NA: neutravidin. **b.** Characteristic fluorescence time trace (grey) of HP1α (3 nM) binding dynamics to a single chromatin fiber, in absence of KAP1. High fluorescence emission reveals the time an HP1α molecule is bound (t_bright_), whereas low fluorescence emission indicates the absence of any bound molecules over a given time (t_dark_). The trace is fitted with a step function (yellow). Each intensity peak represents one binding event. **c.** Fluorescence time trace of HP1α – chromatin binding dynamics, in the presence of 100 nM KAP1. The trace is fitted with a step function (red). **d.** Dissociation kinetics: normalized, cumulative histograms of HP1α dwell times in the absence of KAP1 (yellow) or presence of 100 nM KAP1 (red). Both histograms are fitted with a double exponential function. For fit values, see **Table 1. e.** Association kinetics: normalized cumulative histogram of times between binding events for HP1α alone (yellow) or HP1α in the presence of 100 nM KAP1 (red), fitted with mono-exponential function. For fit values, see **Table 1. f.** Dwell times *τ*_off,1_ and **g.** dwell times *τ*_off,2_, demonstrating that KAP1 stabilizes HP1α-chromatin interactions. Numbers indicate % amplitude. Error bars: standard deviation (s.d.), n = 3 - 10 replicates, *: p<0.05 vs. 0 nM KAP1, student’s t-test. For the fit values, see **Table 1. H.** KAP1 reduces on-rates *k_on_*. Error bars: standard deviation (s.d.), n = 3 - 10 replicates, *: p<0.05 vs. 0 nM KAP1, student’s t-test. For the fit values, see **Table 1. i.** A histogram of observed, normalized fluorescence peak intensities in kinetic traces reports on the oligomeric state of chromatin bound HP1α (yellow). This demonstrates that HP1α exists mostly as monomers at 3 nM concentration. Considering the observed labeling efficiency of 60%, the 8% of all observations correspond to dimers. **j.** At 100 nM, KAP1 stabilizes HP1α dimers (28%) (red). However, no higher oligomers of HP1α are observed. As a comparison, the expected fluorescence intensity distribution for 30% dimers and 20% tetramers (with 60% labeling efficiency) is given in grey.

**Table 1.** smTIRF values. Fit results from smTIRF measurements of HP1α in the presence of the indicated concentrations of KAP1. n denotes the number of independent experiments, each contributing > 100 kinetic traces.* values taken from ref.^77^.

| [KAP1](nM) | $\tau_{off,1}$ (s) | % A<sub>1</sub> | $\tau_{off,2}$(s) | % A<sub>2</sub> | $k_{on}$ (M<sup>-1</sup>s<sup>-1</sup>) x 10<sup>6</sup> | n |
| --- | --- | --- | --- | --- | --- | --- |
| 0* | 0.25 ± 0.03 | 87 ± 7 | 2.26 ± 1.22 | 13 ± 7 | 3.64 ± 1.56 | 10 |
| 50 | 0.29 ± 0.05 | 89 ± 3 | 3.64 ± 0.47 | 11 ± 3 | 1.03 ± 0.37 | 4 |
| 100 | 0.28 ± 0.02 | 85 ± 3 | 5.79 ± 1.03 | 15 ± 3 | 1.42 ± 0.95 | 4 |
| 200 | 0.27 ± 0.04 | 87 ± 2 | 5.47 ± 0.93 | 13 ± 2 | 1.43 ± 0.45 | 3 |
| 400 | 0.23 ± 0.02 | 86 ± 11 | 5.03 ± 0.65 | 14 ± 11 | 0.77 ± 0.44 | 3 |

To more quantitatively elucidate the effect of KAP1 on HP1α binding dynamics we tested the effect of KAP1 concentrations ranging from 50 – 400 nM. While the fast binding times *τ*_off,2_ did not exhibit a systematic dependence on KAP1 concentration (**Figure 6f**), *τ*_off,2_ was increased by around three-fold at KAP1 > 100 nM (**Figure 6g**). Thus, KAP1 stabilizes bivalent chromatin interactions of HP1α dimers, consistent with the fact that a KAP1 dimer can only bind one HP1 dimer. Conversely, binding rate constants (**Figure 6h**) were reduced by KAP1, consistent with slower diffusion dynamics of the HP1α-KAP1 complex. Finally, an analysis of the fluorescence intensities of the single-molecule observations allowed us to quantify the oligomeric state of HP1α in our experiments. In the absence of KAP1, HP1α was mainly monomeric (due to the low concentrations employed) and only 8% dimers were observed (**Figure 6i**). Addition of 100 nM KAP1 resulted in an increase of dimers (up to 28%, **Figure 6j**). Importantly, no higher oligomeric species of HP1α were detected (**Figure 6j**), consistent with our previous observations that one KAP1 dimer can only bind a single HP1α dimer at a time.

Thus, in the chromatin context represented by these conditions and using trimethylated nucleosomes, a KAP1 dimer is able to bind one dimer of HP1 with two binding sites for the histone tails packing the chromatin fiber. These bivalent interactions are longer-lived than the interactions of HP1 alone, such that in the cell, this might translate into a more durable heterochromatin state that is able to spread along the DNA. It remains to be studied how the remaining PTMs and chromatin modifiers influence or are affected by this asymmetric architecture.

## Discussion

Gene expression in response to physiological and environmental stimuli is controlled by the cell at different levels by chromatin chemical modifications such that effector proteins can interpret them and alter the chromatin state^78–80^. These proteins are thought to have multiple roles and work in an integrated fashion, being modulated by PTMs themselves. KAP1 is a master regulator that exemplifies this level of complexity and coordination between PTMs and the binding of protein partners.

KAP1 is not only SUMOylated but also phosphorylated at multiple sites^81^ and the same applies to its binding partners, like HP1^49^. The cross-talk between the PTMs within KAP1 and HP1 in fact affects their function in transcription control and DNA damage response^8^, and it is still not well understood at the molecular level. This is mainly due to the very flexible nature of these proteins, both containing multiple domains linked by disordered linkers, which so far have precluded the solution of their full length structure. Therefore, we have used here an integrative modeling approach combining solution scattering diffraction data, flexible fitting, biophysical and biochemical analyses as well as single-molecule experiments to reveal the molecular architecture of full-length KAP1 and found an unexpected asymmetry for its C-terminal domains. On one hand, as other TRIM family members, KAP1 contains an N-terminal RBCC domain that consistently folds into an antiparallel long coiled-coil dimer, with the RB_1_B_2_ modules located at the opposite ends of it. On the other hand, the C-terminal domains appear to have an unexpected conformation in solution. Flexible fitting to the SAXS scattering curves resulted univocally in an ensemble of conformations featuring one PHD-Br domain close to one RB_1_B_2_ domain, leaving the second not interacting with the main core of KAP1, and more accessible. This solution is reminiscent of previous findings for the C-terminal PRYSPRY domains of TRIM25^55^, where although in crystallographic conditions both domains were interacting along the CC region, in solution, SAXS showed transient interactions with the CC domain, hinting to a much more dynamic conformation for the C-termini in more physiological conditions^55,82^.

Is this native asymmetry in solution associated with KAP1 function? On one hand, auto-SUMOylation assays have shown that the RING domain does play a role in the SUMOylation of the C-terminus. As shown by us and others^37^, SUMOylation happens at lysine residues at the C-terminal end and some SUMOylation sites remain SUMOylated after the PHD-Br domain is compromised with catalytic incompetent mutations (e.g., C651A). The remaining SUMOylation could be due to (i) the existence of another E3 ligase active site or (ii) the PHD being still active despite C651A mutation. Here, we provide evidence that the RING domain can act as an additional catalytic domain SUMOylating those sites or can act synergistically with the PHD domain for the same purpose. Unfortunately, KAP1 bearing RING truncations together with PHD-Br inactive mutations proved to be insoluble, not allowing us to definitively distinguish between these two situations. However, the proximity of the RING and the PHD-Br domains suggested by our SAXS-based asymmetric KAP1 models are consistent with this picture. RING domains in other TRIM family members dimerize to preserve E3 ligase activity (e.g., TRIM32^61^, TRIM5a^58^ or TRIM23^83^), but the RING domain of KAP1 appears to be monomeric (Modis and coworkers, personal communication). This finding supports our model in suggesting a possible synergy between the RING and the PHD domain, which has itself a RING like fold. Therefore, we conclude that KAP1 has two separate SUMO E3 ligase active sites ^31,32,37^, which might work independently or synergistically depending on the substrate available.

One obvious reason for KAP1 dimer asymmetry is to modulate KAP1 interactions with binding partners like HP1 (**Supplementary Figure 8** and ref. ^84^). Previous data using the CSD and a small peptide from KAP1 showed that as expected, the CSD dimerized and bound one KAP1 peptide^46^. According to our MALS data, this interaction is maintained when using FL proteins but involving only one of the two available HP1BDs of KAP1. This unexpected stoichiometry suggests then that the asymmetric structure of KAP1 is regulating HP1 binding accessibility in isolation and in the chromatin context. Within chromatin fibers, KAP1 dimers recruit indeed HP1 as dimers to trimethylated nucleosomes such that two chromo domains per complex can reach for the histone tails. As this interaction is more durable with KAP1, we can speculate that in cells, the binding of KAP1 stabilizes HP1 dimerization in the chromatin facilitating heterochromatin formation and spreading. Regarding the mode of interaction, it is interesting to notice how more than half of the known KAP1 PTMs occur next to the HP1 binding domain. However, the HP1-KAP1 interaction is independent from PTMs, suggesting that PTMs may play a role possibly affecting complex stability and chromatin interactions. Similarly, this functional asymmetry in recruiting partners with controlled stoichiometry can be present for other KAP1 interacting partners engaged by the C-termini, such as SETDB1, or that interact with the CC domain, such as the MAGE (melanoma antigen genes) protein or KRAB-ZFPs^9,26,43^. For the latter in fact it has been recently shown that KAP1 FL binds only one KRAB domain of the ZNF93 at around the CC domain (Modis and coworkers, personal communication).

What is then the ultimate purpose of KAP1 asymmetry? The answer can be linked to the multiple functions of KAP1, such that one RING/PHD-Br unit of the dimer is responsible for SUMOylation activity, while the other PHD-Br domain is SUMOylated and able to interact with histone modifiers. Both units could be independently modulated by PTMs or bind partners allowing for multiple layers of regulation and fast responses. The flexibility and intrinsically disordered nature of the linker regions of the KAP1 dimer would allow for this asymmetry, for an extended line of residues available for PTM regulation^85^ and also for the rapid evolution of the micro-domains within it (like the HP1BD), which could mutate to recruit different binding partners as needed without having to maintain a stable fold^86^.

In conclusion, our findings shed light for the first time on the molecular architecture of full-length KAP1, its interaction with HP1 in the context of chromatin fibers and the implications of its native flexibly and resulting asymmetry. Canonical structural biology techniques still struggle to provide a high-resolution characterization of systems of this kind, where multiple domains are connected by disordered linkers producing natively flexible complexes capable of plastic, dynamic interactions modulated by specific environmental conditions. Studying such dynamic systems using an integrative approach able to combine experimental data and molecular modeling appears to be a powerful resource to understand their behavior in the cellular environment.

## Methods

### Expression and purification of recombinant proteins

Recombinant proteins were expressed and purified from *Escherichia coli (E. coli)*. The KAP1 variants DNA coding sequences were optimized for *E. coli* expression (Genescript) and cloned into the first multi-cloning site (MCSI) of the pETDuet-1 vector, preceded by a His6 tag and a Tobacco Etch Virus (TEV) protease cleavage site. The plasmids were transformed in Rosetta (DE3) and Rosetta (DE3) pLys cells (Promega). Cells were grown to an optical density of 0.7-0.8 in Luria-Bertani (LB) media. Protein expression was induced by the addition of 1 mM isopropyl β-D-1-thiogalactopyranoside (IPTG) and subsequent growth over night at 20°C. Cell pellets were resuspended in lysis buffer (20 mM HEPES, pH 7.5, 500 mM NaCl, 10% (v/v) glycerol, 2 mM Tris(2-carboxyethyl)phosphine (TCEP), 5 mM Imidazole and cOmplete™ Protease Inhibitor Cocktail (Roche)) and then lysed using high pressure homogenizer (Avestin Emulsiflex C3). The resulting suspensions were centrifuged (13000 rpm for 35 min at 4°C) and the supernatants were applied to an HisTrap HP column (GE Healthcare) previously equilibrated with loading buffer (20 mM HEPES, pH 7.5, 10% (v/v) glycerol, 500 mM NaCl). The proteins were eluted with a gradient over 40 column volumes of elution buffer (20 mM HEPES, pH 7.5, 500 mM NaCl, 10% (v/v) glycerol, 500 mM Imidazole). Subsequently, pure fractions were buffer exchanged into final buffer (20 mM HEPES, pH 7.5, 500 mM NaCl, 10% (v/v) glycerol, 2 mM TCEP) using a HiPrep Desalting column (GE Healthcare) and stored over night at 4°C in final buffer. The proteins were additionally purified by Size Exclusion Chromatography (SEC) in final buffer, flash frozen in liquid nitrogen and stored at - 20°C. HP1α was purified as previously described^77^.

### Size-exclusion chromatography coupled to multi-angle light scattering

The molecular weights of the constructs were determined by size exclusion chromatography coupled to a multi angle light scattering detector (SEC-MALS). The mass measurements were performed on a Dionex UltiMate3000 HPLC system equipped with a 3 angles miniDAWN TREOS static light scattering detector (Wyatt Technology). The sample volumes of 100 μl at a concentration of 40 μM, were applied to a Superose 6 10/300 GL column (GE Healthcare) previously equilibrated with 20 mM HEPES pH 7.5, 300 mM NaCl, 2 mM TCEP at a flow rate of 0.5 ml/min. The data were analyzed using the ASTRA 6.1 software package (Wyatt technology), using the refractive index of the buffer as a baseline and the refractive index increment for protein dn/dc = 0.185 ml/g.

### Analytical Ultra Centrifugation

Sedimentation velocity data of the N-terminal RBCC domain of KAP1 (0.5 mg/mL) were obtained using a Beckman Optima XL-I analytical ultracentrifuge (Beckman Coulter) with an absorbance optical system. Epon double-sector centerpieces containing 200 μl of protein solution and 300 μl of sample buffer were centrifuged at 55000 rpm and 20°C and data acquired with a radial increment of 0.003 cm with no delay between scans. The sedimentation of the protein was monitored at 292 nm and 280 nm. Data analysis was performed with Sedfit^87^. Buffer viscosity (0.01002 cp), density (1.04 g/cm3) and protein partial specific volumes (0.72 g/cm) were estimated using the program Sednterp^88^.

### Small-Angle X-Ray Scattering data collection and analysis

To exclude sample aggregation, the proteins were analyzed by size-exclusion chromatography in line with small-angle X-Ray scattering (SEC-SAXS). The data were collected at the European Synchrotron Radiation Facility (ESRF), beamline 29 (BM29) at a wavelength of 0.99 Å with a sample to detector distance of 2.867 m and a PILATUS 1M detector, covering a momentum transfer of 0.0025 < q > 0.6 Å^−1^ [q = 4πsin (θ)/λ]. Measurements were made at 18°C. The samples were applied to a Superose 6 10/300 GL column (GE Healthcare) at a concentration of 9-12 mg/ml and run at a flow rate of 0.75 ml/min in 20 mM HEPES, pH 7.5, 500 mM NaCl, 10% (v/v) glycerol, 2 mM TCEP. During the elution, 2160 scattering measurements were taken with 1s time-frames. The in-house software *BsxCuBE* (Biosaxs Customized Beamline Environment) connected to a data processing pipeline (EDNA)^89^ was used to control the real time data display (two dimensional and one dimensional) and to provide the first automatic data processing up to a preliminary *ab initio* model. SAXS data were analyzed using the ATSAS package version 2.8.3^90^ and ScÅtter^91^. For each sample, using Chromixs^92^, an elution profile was generated with the integrated intensities plotted versus the recorded frame number. Using Chromixs, 30 buffer frames were averaged and used to (i) subtract the buffer average from each frame of the sample peak selected and (ii) calculate the corresponding Radius of Gyration (*R*_g_). The subtracted peak region was selected in Chromixs and averaged to generate the final scattering curve used for subsequent analysis. The scattering curves were initially viewed in *PRIMUS*^93^ where the *R*_g_ was obtained from the slope of the Guinier plot within the region defined by *q*_min_ < *q* < *q*_max_ where *q*_max_ < 1.3/*R*_g_ and *q*_min_ is the lowest angle data point included by the program. When one dimension of a scattering particle is greater than the other two (e.g. a rod particle), “Rod” Guinier analysis in PRIMUS can be used to calculate the radius of gyration of its cross-section, R_gc_ (**Supplementary Figure 3**). The *P*(*r*) function, the distribution of the intra-atomic distances (r) in the particle, was generated using the indirect transform program *GNOM*^94^. The maximum distance (*D*_max_) was selected by letting the *P*(*r*) curve decay smoothly to zero. As our molecules are rod-like and flexible, the *R*_g_ was also estimated from the *P*(*r*) function, such that, unlike the Guinier *R*_g_ estimation, the *P*(*r*) R_g_ calculation takes the whole scattering curve into account. *Ab initio* models were produced using GASBOR^64^ imposing a P1 symmetry with prolate (elongated protein) anisotropy until a χ^2^ of 1 was reached. DATPOROD, DATMOW and DATVC within the ATSAS package were used to estimate the Porod Volume (Vp) and the concentration-independent estimate of the MW for the proteins. The final figures were generated using VMD^95^, PyMOL (Schrödinger, LLC) and Chimera^96^.

### Molecular modelling and flexible SAXS data fitting

The preliminary KAP1 model was created using existing, homologous structures of the RING, B-box and coiled-coil α-helical domains, and the solved NMR structure of the PHD-Br domain (PDB: 2RO1)^40^. Specifically, the SWISS Model server^97^ was used to construct the RBCC domain using the following templates: the RING domain was based on the dimer of Rad18 (PDB: 2Y43)^98^, the B-box1 was based on the B-box domain of MuRF1 (PDB: 3DDT)^99^, the B-box 2 was based on the B-box domain of TRIM54 (PDB: 3Q1D), and the coiled-coil domain was based on the coiled-coil of TRIM69 (PDB: 4NQJ)^60^. To model the TSS domain in the central part of KAP1 we used the Robetta server^100^, while MODELLER v9.14^101^ was used for the HP1-binding domain, assigning as a template the structure of EMSY protein in complex with HP1 (PDB: 2FMM)^102^ and the structure of the small subunit of the mammalian mitoribosome (PDB: 5AJ3)^103^. Finally all the missing loops were modeled by Rosetta loop modeling application v3.5^104^. In the process of generating starting models for fitting to SAXS data, each PHD-Br domain was randomly rotated and translated at a maximum distance of 140 Å (distance D in **Figure 3b**) from the RBCC domains using VMD^95^. Afterwards, the linker region between each PHD-Br domain and its respective RBCC domain was modelled as a random coil using MODELLER^101^. 1000 models were generated and flexibly fitted to the SAXS data through a χ^2^–minimizing optimization procedure based on the nonlinear Cartesian NMA method called NOLB^66^ and a novel SAXS profile calculator called Pepsi-SAXS^65^. More precisely, for each initial model we performed 100 optimization iterations. Each iteration comprised the computation of 60 slowest normal modes (using the NOLB tool), nonlinear structure deformation along these modes, and choosing the deformation with the least χ^2^ value to the experimental scattering profile (using Pepsi-SAXS). A steepest-descent minimization algorithm was used at the end of each iteration to keep the local topology (bonds and angles) in agreement with the initial structure. The flexible fitting method is available as a standalone executable called Pepsi-SAXS-NMA at (https://team.inria.fr/nano-d/software/pepsi-saxs/). The choice of 60 normal modes was based on the observation of high flexibility of the linker region. This number of modes allowed us to explore plenty of plausible conformations of the linker. The chosen slowest modes did not change the structure of the rigid domains, but only changed their relative orientation. ~60% of the initial models converged to low values of χ^2^ within the wall-time assigned for fitting (24h). The resulting structures produced a statistically relevant ensemble and were clustered using a new method tailored for clusters exhibiting Gaussian distributions, typical of structural ensembles. As for collective variables, the distance between the center of mass of the PHD-Br domain and that of the closest RB_1_B_2_ module was computed for each protomer. The models were classified according to these two distances and the centroids were extracted (**Figure 3d**).

### Negative Stain Electron Microscopy

KAP1 Fl was diluted to 0.05 mg/ml in 20 mM HEPES pH 7.5, 300 mM NaCl and 2 mM TCEP and crosslinked with 0.1% glutaraldehyde for 2h at 23°C. The reaction was stopped by addition of 100 mM Tris-HCl pH 7.5. The sample was diluted 20 times, and adsorbed to a glow-discharged carbon-coated copper grid (EMS, Hatfield, PA, USA) washed with deionized water and stained with a solution of 2% uranyl acetate. The grids were observed using an F20 electron microscope (Thermo Fisher, Hillsboro, USA) operated at 200 kV. Digital images were collected using a direct detector camera Falcon III (Thermo Fisher, Hillsboro, USA) with 4098 X 4098 pixels. The magnification of work was 29000X (px=0.35 nm), using a defocus range of −1.5 μm to −2.5 μm. After manual picking of 400 particles, Relion^105^ was used to sort them into 2D class averages.

### H3K9me3 synthesis

H3K9me3 was synthesized as previously described^77^. In short, the peptide H3(1–14) K9me3-NHNH_2_ (carrying a C-terminal hydrazide) was synthesized by solid phase peptide synthesis (SPPS). The truncated protein H3(Δ1–14)A15C was recombinantly expressed as N-terminal fusion to small ubiquitin like modifier (SUMO), the N-terminal SUMO was cleaved by SUMO protease and the protein purified by RP-HPLC. For ligation, in a typical reaction 3 μmol H3(1–14)K9me3-NHNH_2_ was dissolved in ligation buffer (200 mM phosphate pH 3, 6 M GdmCl) at −10°C. NaNO_2_ was added dropwise to a final concentration of 15 mM and incubated at −20°C for 20 min. H3(Δ1–14)A15C was dissolved in mercaptophenyl acetic acid (MPAA) ligation buffer (200 mM phosphate pH 8, 6 M GdmCl, 300 mM MPAA) and added to the peptide, followed by adjustment of the pH to 7.5. The ligation was left to proceed until completion (as observed by RP-HPLC). The product (H3K9me3A15C) was purified by semi-preparative RP-HPLC. For desulfurization, H3K9me3A15C was dissolved in tris(2-carboxyethyl)phosphine (TCEP) desulfurization buffer (200 mM phosphate pH 6.5, 6 M GdmCl, 250 mM TCEP). Glutathione (40 mM) and a radical initiator, VA-044 (20 mM), were added, followed by a readjustment of the pH to 6.5. The reaction mixture was incubated at 42°C until completion (verified by RP-HPLC and ESI-MS). The final product, H3K9me3, was purified by semi-preparative RP-HPLC, lyophilized and kept at −20°C for further use. A final characterization of the modified histones was done by analytical RP-HPLC and ESI-MS.

### Chromatin preparation

As previously described^77^, chromatin arrays were reconstituted at a concentration of around 1 μM per mononucleosome, at a scale of 100 pmol. Array DNA (12×601 with 30 bp linker DNA) was mixed with 1 equivalent (per nucleosome positioning sequence) of recombinant human histone octamer, containing H3K9me3, in reconstitution buffer (10 mM Tris pH 7.5, 10 mM KCl, 0.1 mM EDTA) containing 2 M NaCl. 0.5 molar equivalents of MMTV DNA was added to prevent oversaturation. In the case of reconstituted chromatin containing linker histone, 0.5, 1 or 1.5 equivalents H1.1 were also added to the DNA/octamer mixture. The reactions were gradually dialyzed from high salt buffer (10 mM Tris pH 7.5, 1.4 M KCl, 0.1 mM EDTA) to reconstitution buffer over 12 h with a two-channel peristaltic pump. After the dialysis the reconstituted chromatin arrays were analyzed by non-denaturing 5 % polyacrylamide gel electrophoresis (PAGE) in 0.5 x Tris-Borate-EDTA (TBE) running buffer or on a 0.6 % agarose gel following ScaI restriction digest.

### HP1 labeling

HP1α was labeled as described in ref.^77^. A short tripeptide (Thz-G_2_-C_3_-CONH_2_, Thz: thiazolidine) was synthesized by SPPS. 1 mg Atto532-iodoacetamide (5 eq.) was used to label the tripeptide in 200 mM phosphate pH 7.3, 5 M GdmCl. The reaction was followed by analytical-HPLC and MS, after completion quenched by addition of TCEP, purified by semi-preparative RP-HPLC, and lyophilized. The thiazolidine was opened by treatment with 2 M methoxylamine at pH 5. The labeled tripeptide was finally purified by semipreprative RP-HPLC, lyophilized and kept at −20°C until further use. HP1α was expressed as a fusion to the *Npu^N^* split-intein at its C-terminus, followed by a hexahistidine tag. Expression was induced in *E. coli* BL21 DE3 cells with 0.25 mM IPTG overnight at 18° C. After cell lysis in lysis buffer (25 mM phosphate pH 7.8, 50 mM NaCl, 5 mM imidazole and 1x protease inhibitor/50 mL), HP1α was purified over Ni-affinity resin and eluted with elution buffer (25 mM phosphate pH 7.8, 50 mM NaCl, 400 mM imidazole). Eluted fractions were pooled and further purified by anion exchange chromatography, using a 1 ml HiTrap Q FF column and a gradient from low (25 mM phosphate pH 7.8, 50 mM NaCl) to high salt buffer (25 mM phosphate pH 7.8, 1 M NaCl). A total of 500 μl of the expressed HP1-Npu^N^ fusion constructs at a concentration of 50–100 μM were applied to a column of 125 μl SulfoLink resin slurry, containing an immobilized Npu^C^ peptide^106^. After 5 min incubation, the column was drained, followed by washes with wash buffer (100 mM phosphate pH 7.2, 1 mM EDTA, 1 mM TCEP) containing high (500 mM NaCl), intermediate (300 mM NaCl) and low salt (150 mM NaCl). 1 mM of tripeptide in labeling buffer (100 mM phosphate pH 7.8, 50 mM MPAA, 200 mM MESNA, 150 mM NaCl, 10 mM TCEP, 1 mM EDTA) was added to the column and incubated for 16 h at room temperature. The column was drained and the eluate was collected. The column was further washed using elution buffer (100 mM phosphate pH 7.2, 200 mM MESNA, 150 mM NaCl, 10 mM TCEP, 1 mM EDTA). The eluted protein was finally purified by size exclusion chromatography with a superdex S200 10/300 GL in gel filtration buffer (50 mM Tris pH 7.2, 150 mM NaCl, 1 mM DTT. Fractions containing purified labeled HP1 were pooled and concentrated, glycerol was added to 30 % (v/v), frozen aliquots were stored at −80°C. A final characterization of the purified and labeled HP1 proteins was done by analytical RP-HPLC and ESI-MS.

### Single-molecule assays

Single-molecule measurements were performed as previously reported^77^. Glass coverslips (40 × 24 mm) and microscopy slides (76 × 26 mm) containing four drilled holes on each side were cleaned by sonication in 10 % alconox, acetone and ethanol with washing in miliQ H_2_O between each step. The slides/coverslips were incubated for 1 h in a mixture of concentrated sulfuric acid to 30 % hydrogen peroxide (3:1). The coverslips and slides were thoroughly washed with miliQ H_2_O, sonicated in acetone for 15 min and then submerged in acetone containing 2 % (3-aminopropyl)triethoxysilane (APTES) for silanization. The slides and coverslips were dried with a nitrogen flow and strips of double-sided tape were sandwiched between a coverslip and a slide to create four channels. The glass coverslips were passivated with a solution of 100 mg/ml mPEG(5000)-succinimidyl carbonate containing 1 % biotin-mPEG-succinimidyl carbonate for 3 h. The channels were subsequently washed with water and T50 buffer (10 mM Tris, 50 mM KCl). For chromatin immobilization, 0.2 mg/ml neutravidin in T50 injected for 5 min, followed by extensive washes with T50 buffer. Then, 500 pM reconstituted chromatin arrays in T50 buffer were injected into the neutravidin treated flow chamber for 5 min, followed by T50 washes and imaging buffer (50 mM HEPES, 130 mM KCl, 10 % (v/v) glycerol, 2 mM 6-hydroxy-2,5,7,8-tetramethylchromane-2-carboxylic acid (Trolox), 0.005 % Tween-20, 3.2 % glucose, glucose oxidase/catalase enzymatic oxygen removal system). Chromatin coverage was observed with a TIRF microscope (Nikon Ti-E) by fluorescent emission in the far-red channel upon excitation by a 640 nm laser (Coherent Obis). Dynamic experiments were initiated by influx of 3 nM Atto532-labeled HP1, as well as the indicated KAP1 concentrations in imaging buffer. All smTIRF experiments were performed at room temperature (22°C). HP1 dynamics were observed with an EMCCD camera (Andor iXon) in the yellow/orange channel using a 530 nm laser line for excitation at 20 W/cm^2^. 10k frames were acquired at 20 Hz over a 25 × 50 μm observation area at a resolution of 160 nm/pixel. Every 200 frames, an image of the chromatin positions in the far-red channel was recorded for drift correction. For each chromatin fiber, an individual trace was extracted using a custom-made semi-automated Matlab (Mathworks) script, as described in ref.^77^. After an initial baseline correction and a drift correction, a peak-finding algorithm was employed to detect individual chromatin array positions. Fluorescence intensity traces for each chromatin position were obtained by integrating over a circle of 2-pixel radius. Individual HP1 fluorescence peaks were included based on point-spread-function (PSF) and distance cut-offs. To ensure that only single-molecules were analysed, peaks exhibiting step-wise bleaching kinetics were excluded from the analysis. Kinetics were extracted from fluorescence traces using a semi-automated thresholding algorithm. Cumulative histograms were constructed from dark and bright intervals and fitted to mono or bi-exponential functions. For intensity analysis, normalized intensity histograms were constructed over several hundred kinetic traces.

### In vitro SUMOylation assay

The in vitro sumoylation assay was conducted using a commercial kit from Abcam (ab139470) using 1μM target protein. The mixtures (E1, E2 ubc9, SUMO1, ATP-Mg, target proteins) were set up in 20 μl of 10X SUMO reaction buffer and incubated at 37°C for 2h. The reactions were collected every 20 min to study the progression of the SUMOylation process. The reaction mixtures were subsequently separated using sodium dodecyl sulfate–polyacrylamide gel electrophoresis SDS-PAGE (4-12%) and were subjected to western blot analysis using anti SUMO1 antibody. Gels were scanned with a Fusion FX 7 (Witec) and analysed using ImageJ^75^. Experiments were performed in triplicate and representative images are shown, while un-cropped gels are shown in **Supplementary Figure 7**.

### Isothermal titration calorimetry

ITC experiments were performed using a MicroCal PEAQ ITC from Malvern. KAP1 FL and HP1 FL were buffer exchanged into 20 mM HEPES pH 7.5, 300 mM NaCl, 2 mM TCEP and concentrated to 1.5 and 5 mg/ml (15 and 200 μM), respectively. HP1 FL containing solution was injected into the KAP1 solution (2 μl of HP1 per injection at an interval of 180 sec, a total of 19 injections into the cell volume of 300 μl, with stirring speed of 800 rpm, at 25° C). HP1 FL was also injected into buffer to determine the unspecific heat of dilution. However, subtracting this experimental heat of dilution was not sufficient to get a good fit, so the last injections were used to better estimate the heat of dilution and subtract a straight line from our data. The experimental data were fitted to a theoretical titration curve (“one set of sites” model) using software supplied by Microcal, with n (number of binding sites per monomer), ΔH (binding enthalpy, kcal/mol) and K_d_ (dissociation constant, M), as adjustable parameters. Thermodynamic parameters were calculated from the Gibbs free energy equation, ΔG =ΔH – TΔS =−RT lnK_a_, where K_a_ is the association constant, and ΔG, ΔH and ΔS are the changes in free energy, enthalpy, and entropy of binding, respectively. T is the absolute temperature, and R = 1.98 cal mol^−1^ K^−1^.

### Data availability

SAXS data and models were deposited in the Small Angle Scattering Biological Data Bank SASBDB with accession code SASDEV6, SASDER7 and SASDEW6. Other data are available from the corresponding authors upon request.

## Acknowledgements

We thank the EPFL Protein Production and Structure Core Facility for providing the equipment for the biophysical characterization of the protein complexes; Natacha Olieric and Bruno Correia for the use of their MALS machines; Cy Jeffries for his assistance with SAXS data preparation and analysis; Martin Moncrieffe for his help with AUC data analysis; Luciano Abriata and Florence Pojer for helpful discussions; BM29 staff scientists for their assistance in SAXS data collection; EPFL for funding.

## Author Contributions

G.F., M.J.M. and M.D.P. designed the study, wrote the manuscript, analysed and interpreted the data. G.F. performed all experiments (apart smTIRF). M.J.M. performed SAXS experiments. G.F., S.T., A.S.K. and S.G. performed integrative modelling. L.C.B and B.F. performed single-molecule experiments and analysis. P.J.L.H. and D.T. supported SUMOylation experiments. D.D. supported TEM experiments. M.T. supported SAXS experiments and analysis. All the authors contributed to edit and review the manuscript. M.J.M. and M.D.P. supervised the study. M.D.P. managed the project.

## Supplementary Information

**Supplementary Figure 1.**
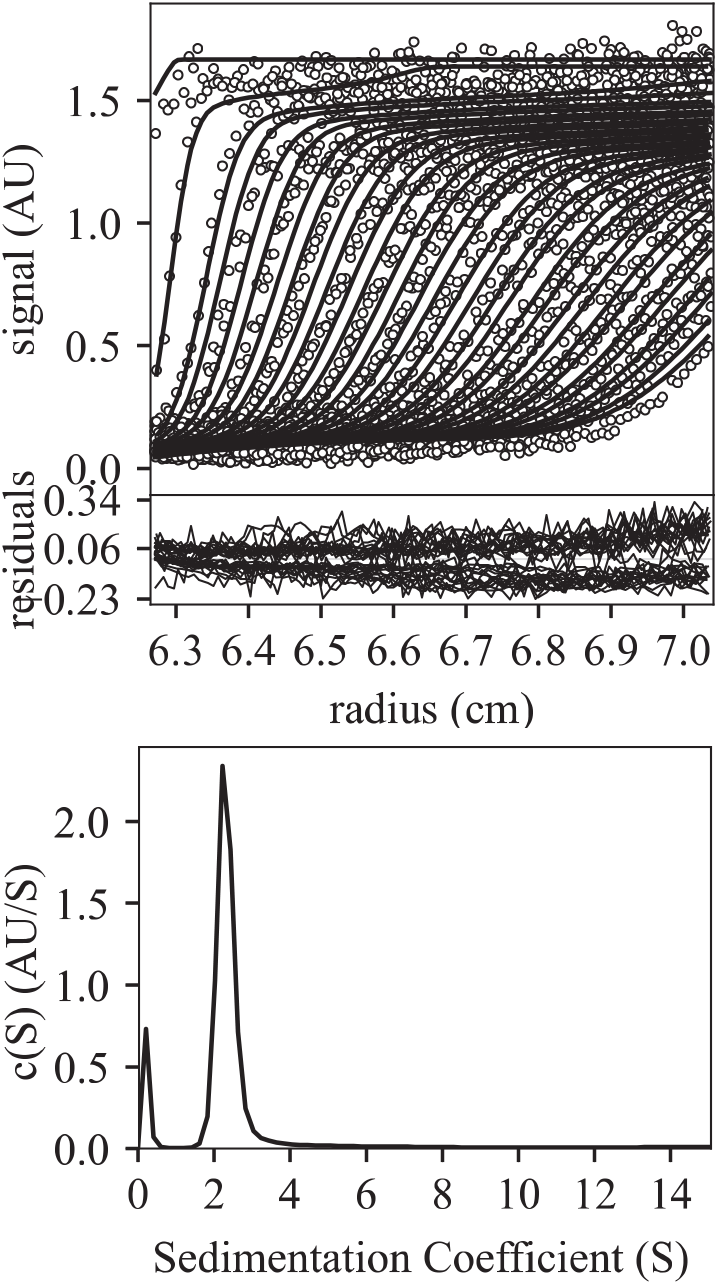
Analytical ultracentrifugation analysis of the KAP1 RBCC domain. The top panel shows the raw sedimentation velocity data, shown as fringe displacements (which is proportional to the protein concentration) plotted against the distance from the axis of rotation. The center panel shows the residuals after fitting to a continuous concentration vs molecular mass, c(M), model. The rmsd for the fit was 0.0546, and the sedimentation coefficient was 2.3 S. The bottom panel shows the distribution of molecular masses obtained from the sedimentation velocity data. Data analysis was performed using the program Sedfit^1^.

**Supplementary Figure 2.**
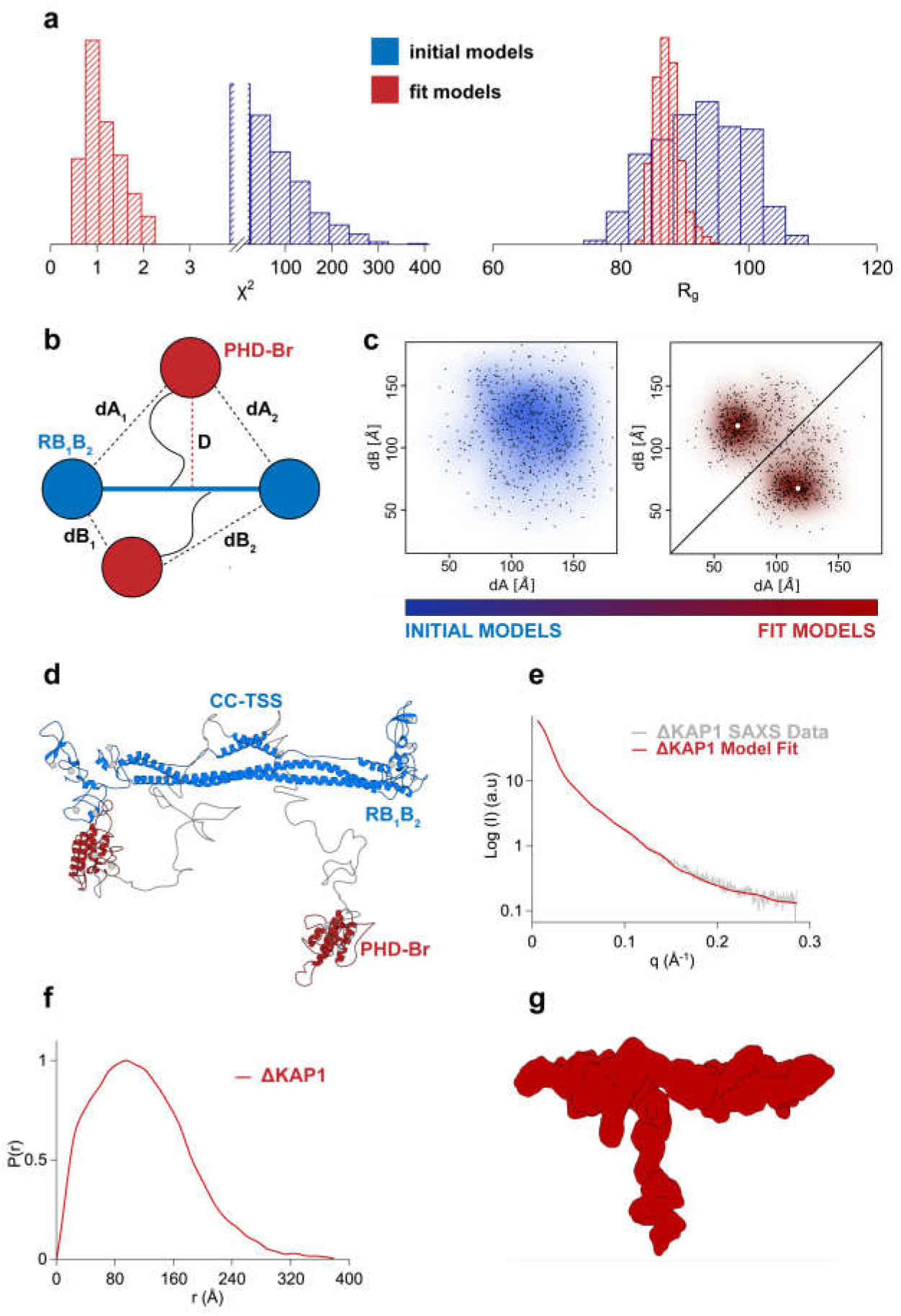
Integrative modelling analysis of ΔKAP1. (**a**) χ plots (left) and *R_g_* plots (right) of the initial (blue) vs. final (red) model structures, with narrower distributions centred on χ =1 and the experimental *R_g_* (See **Supplementary Table 1**). **b.** Schematic representation of the KAP1 dimer where the distances D, dA_1/2_ and dB_1/2_ are displayed. (**c**) Plot of the initial randomized models versus the final fit models according to their dA vs dB distance values. The centroids of the clusters are highlighted by white dots. (**d**) Cartoon representation of the centroid model of KAP1 with the most probable domain arrangement. (**e**) Comparison between the experimental SAXS scattering profile (grey) and the calculated scattering profile (red) from the centroid structure shown in (**d**) using Pepsi-SAXS (χ^2^=0.9). (**f**) Pair distance distribution function (*P*(*r*)) calculated from the SAXS scattering profile. (**g**) GASBOR *ab initio* bead model of ΔKAP1 shown in surface representation.

**Supplementary Figure 3:**
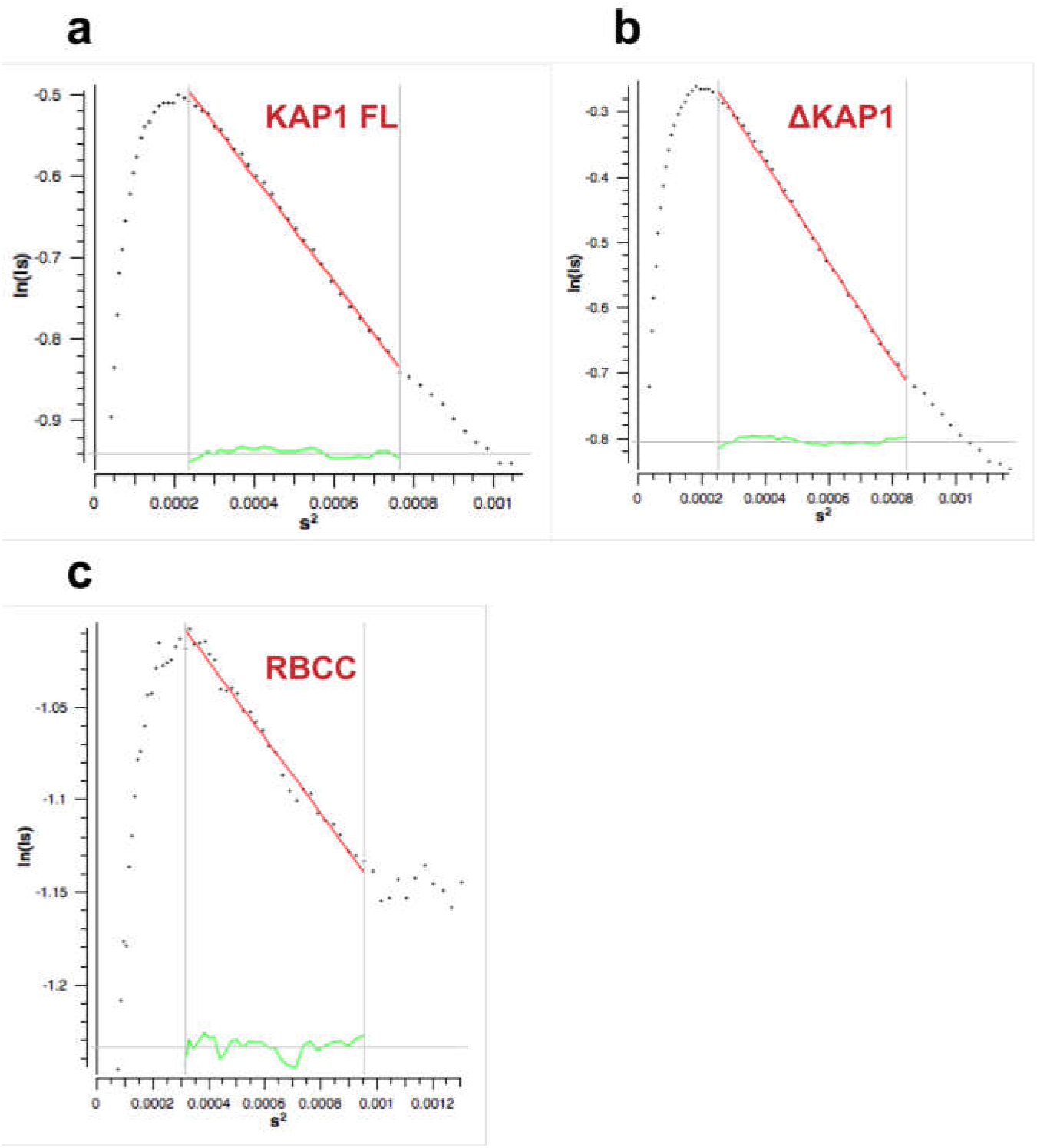
Cross-section R_g_ (R_gc_) plots show that the KAP1 variants are elongated rod-like molecules. Values are shown in **Supplementary Table 1** and were calculated using Primus^2^. s = q = 4πsin (θ)/λ, where θ is the scattering angle and λ is the wavelength of the x-rays.

**Supplementary Figure 4:**
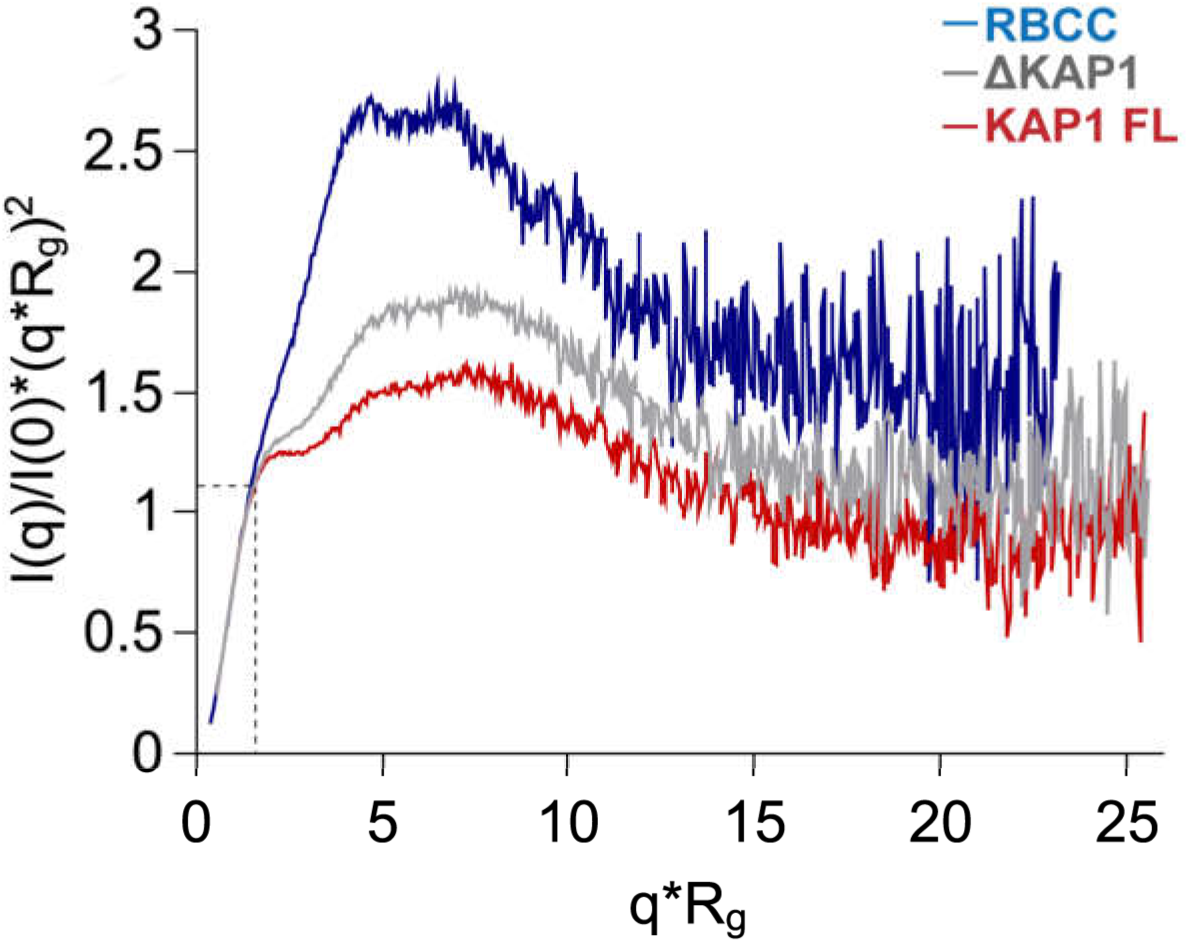
Kratky plot of the KAP1 variants. Dimensionless Kratky Plots produced with ScÅtter^3^ showing that all KAP1 constructs are flexible multi-domain proteins. In dimensionless Kratky Plots, globular proteins always have a maximum at q*R_g_ =3^1/2^ and (q*R_g_)^2^I(q)/I(0)=1.104, which is shown by the dotted lines.

**Supplementary Figure 5:**
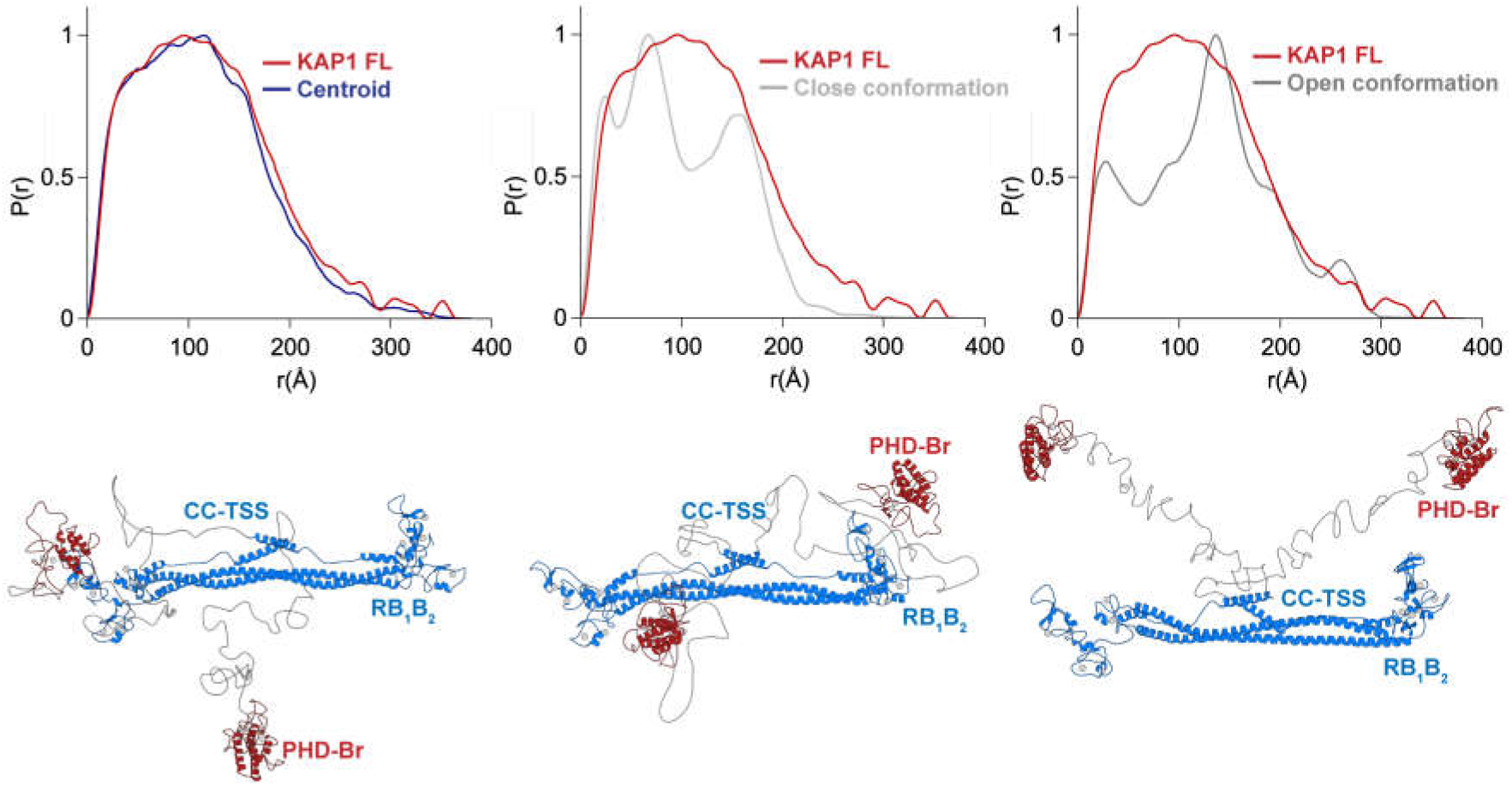
Comparison of *P*(*r*) functions. Superposition of experimental (red) and calculated (blue/grey) *P*(*r*) functions for models of KAP1 FL where the PHD-Br domains are in different positions, as follows: (left) as in the centroid structure, (middle) both close to the RBCC domain and (right) both away from the RBCC domain. The *P*(*r*) functions were calculated using ScÅtter^3^.

**Supplementary Figure 6:**
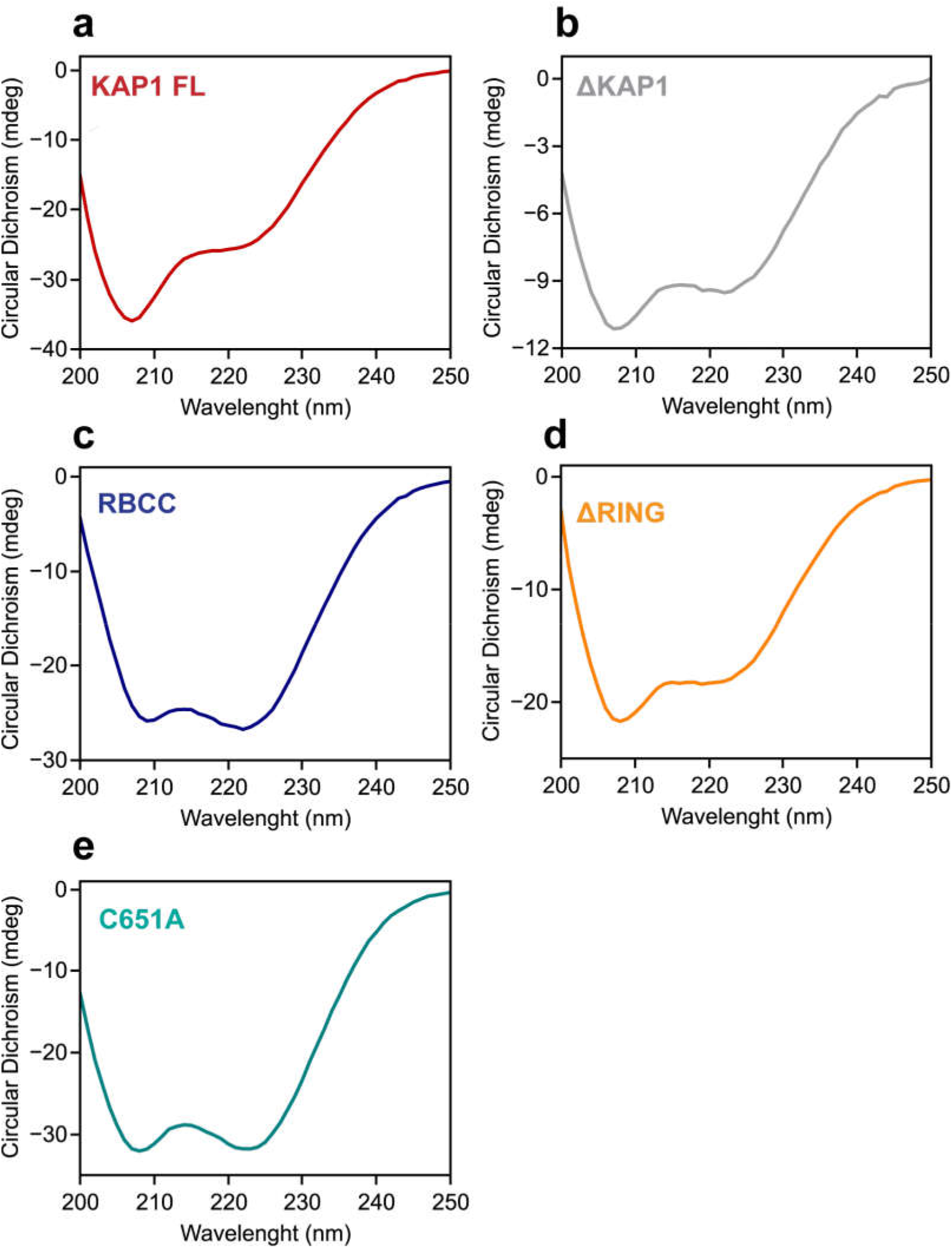
Circular Dichroism (CD) analysis of the KAP1 constructs used in this study shows that they are folded. CD data were collected using a Chirascan CD Spectrometer and a 0.1 cm quartz cell. Sample concentration ranged from 0.2 mg/ml to 0.8 mg/ml in Phosphate Buffer (1x PBS). Scans were measured from 200 to 250 nm, under continuous scanning mode with a 1 nm data pitch, a scan speed of 50 nm/min and a response time of 1 s. The figure shows the average of 3 spectra, which are buffer subtracted.

**Supplementary Figure 7:**
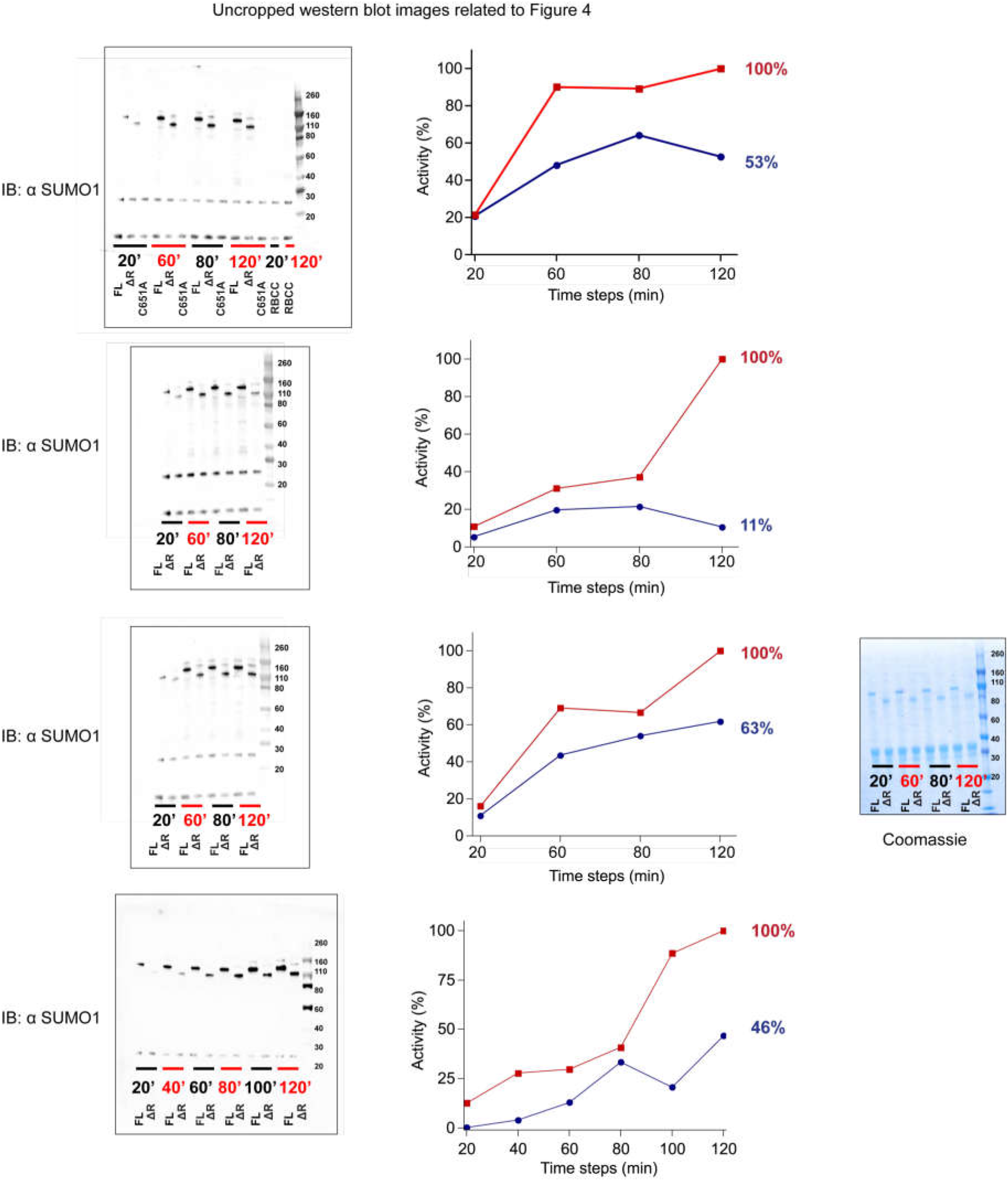
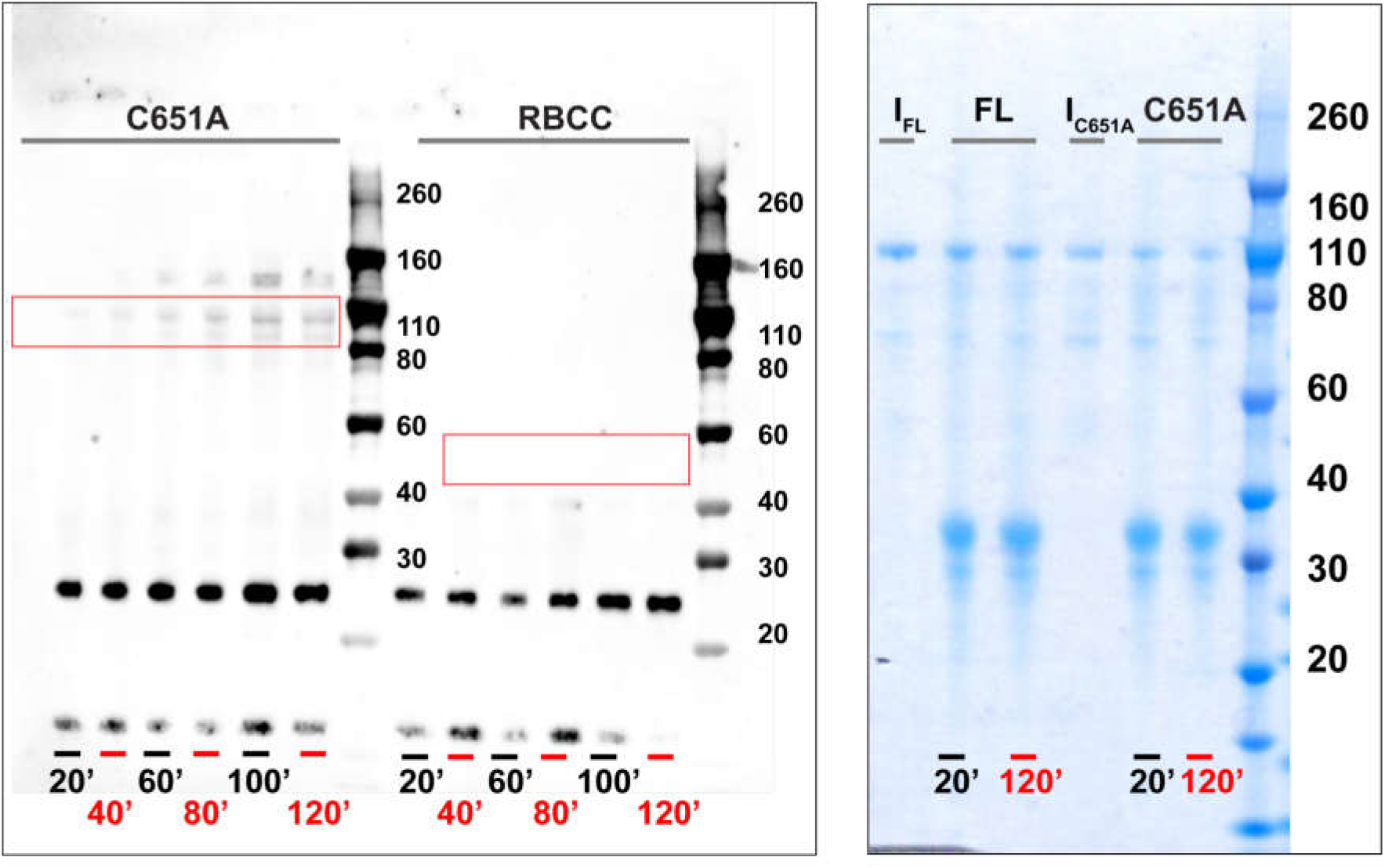
Un-cropped western blots related to Figure 4. The first four panels show the experimental replicas for the auto-SUMOylation assay comparing KAP1 FL and ΔRING. The Coomassie stained SDS-PAGE is shown on the right as loading control. The bottom panel shows the auto-SUMOylation assay using the C651A mutant and KAP1 RBCC. As expected, KAP1 RBCC did not get SUMOylated. When comparing the MW shifts of the proteins upon SUMOylation, it is interesting to notice how when the PHD is active, the size change is large (50 kDa, seen for FL and ΔRING) but when the PHD is compromised, the size change is very small (seen for the C651A mutant). On the right, the Coomassie stained SDS-PAGE of the FL and C651A reactions is shown as loading control, where non SUMOylated proteins are indicated by I FL and I C651A.

**Supplementary Figure 8:**
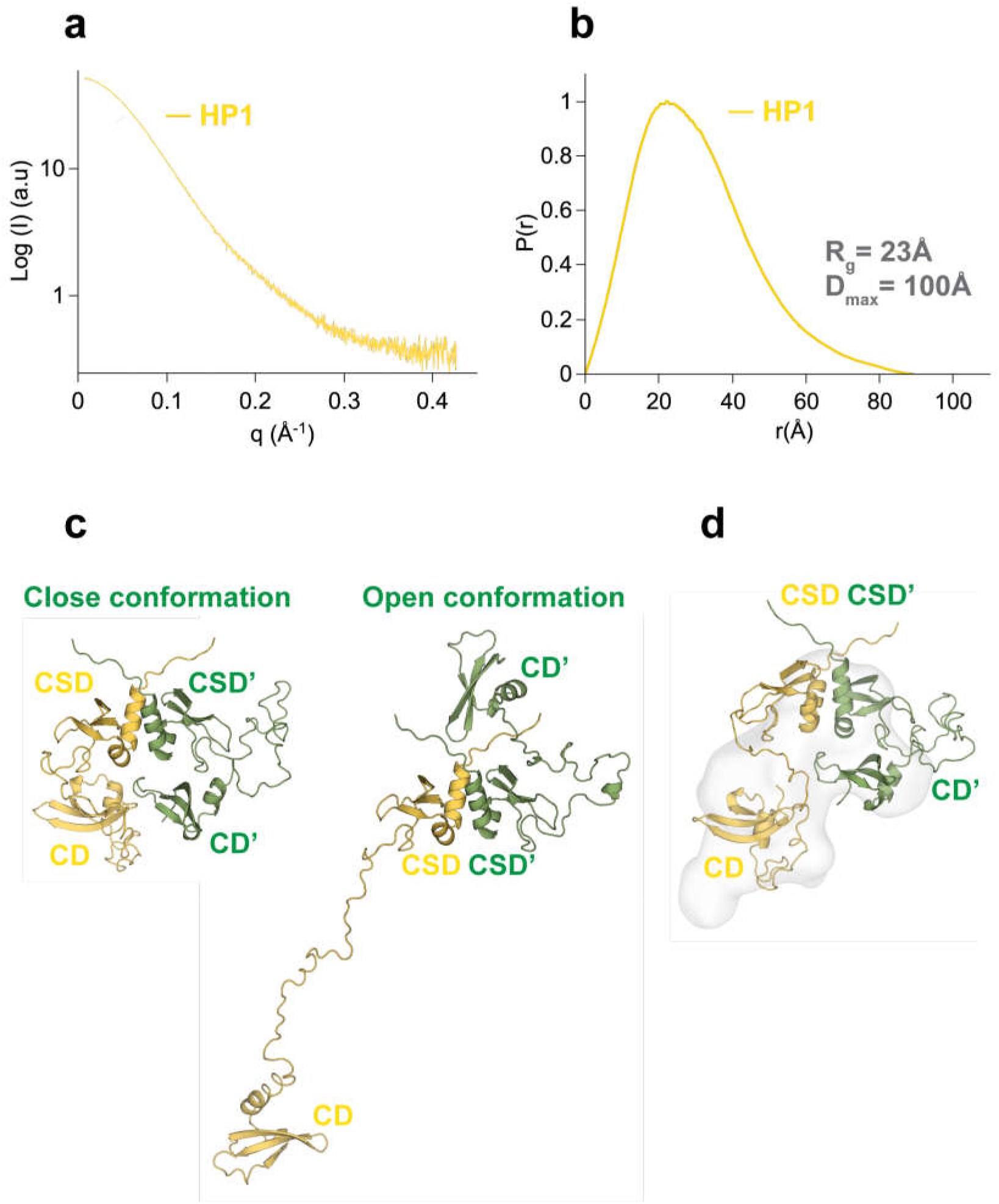
Analysis of the conformation of HP1 FL using SAXS. (**a**) SAXS scattering profile. (**b**) Pair distance distribution function, *P*(*r*). The insert shows the resulting *R_g_* and *D_max_* values. (**c**) The HP1α FL model was built using PDB files 1S4Z for the CSD and 1Q3L for the CD and letting EOM (ensemble optimization method)^4^ build the flexible linker to create ensembles of conformations of the dimer. EOM ensembles contained only 2 structures: a closed one (left) and an open one (right). (**d**) Closed HP1α model superimposed in the *ab initio* shape created directly from the SAXS data using GASBOR.

**Supplementary Table 1:** SAXS parameters.

| | KAP1 FL | ΔKAP1 | RBCC |
| --- | --- | --- | --- |
| **SASPDB accession code** | SASDEV6 | SASDER7 | SASDEW6 |
| **Data collection** |  |  |  |
| Instrument |  | ESRF BM29 |  |
| Beam size at sample ($\mu\text{m}$) | | 700x700 | |
| Wavelength ($\text{\AA}$) | | 0.992 | |
| q range ($\text{\AA}^{-1}$) | | 0.025-0.500 | |
| Detector |  | Pilatus 1M |  |
| Detector distance (m) |  | 2.867 |  |
| Exposure (s per image) |  | 1 |  |
| Column |  | Superose 6 Increase 10/300 |  |
| Flow rate (ml/min) |  | 0.5 |  |
| Sample volume ($\mu\text{l}$) | | 100 | |
| Sample concentration (mg/ml) | 15 | 12 | 12 |
| Temperature (K) |  | 293 |  |
| **Structural parameters** |  |  |  |
| $R_g$ ($\text{\AA}$) Guinier | 90 | 89 | 83 |
| $R_g$ ($\text{\AA}$) P(r) | 95 | 95 | 89 |
| $R_{gc}$ ($\text{\AA}$) | 35.8 | 38.8 | 20.2 |
| $D_{max}$ ($\text{\AA}$) | 380 | 380 | 370 |
| Porod volume ($\text{\AA}^3$) | 710000 | 853000 | 381000 |
| **Molecular mass determination** |  |  |  |
| Theoretical MW (kDa) | 183 | 175 | 92 |
| MALLS MW (kDa) | 190 | 183 | 88 |
| DATPOROD MW (kDa) | 444 | 533 | 238 |
| DATVC MW (kDa) | 255 | 290 | 117 |
| DATMOW MW (kDa) | 247 | 321 | 145 |
| **Data analysis software** |  |  |  |
| Data reduction |  | PRIMUS & ScÅtter |  |
| *Ab initio* modelling |  | GASBOR |  |
| Homology modelling |  | Swiss Model/I-tasser |  |
| Computation of model fitting to data |  | Pepsi-SAXS |  |
| 3D graphics representation |  | Pymol |  |

## References

1. Cheng, C.-T., Kuo, C.-Y. & Ann, D. K. KAPtain in charge of multiple missions: Emerging roles of KAP1. World J. Biol. Chem. 5, 308–320 (2014).

2. Iyengar, S. & Farnham, P. J. KAP1 Protein: An Enigmatic Master Regulator of the Genome. J. Biol. Chem. 286, 26267–26276 (2011).

3. Cammas, F. et al. Mice lacking the transcriptional corepressor TIF1beta are defective in early postimplantation development. Dev. Camb. Engl. 127, 2955–2963 (2000).

4. Cammas, F. et al. Cell differentiation induces TIF1beta association with centromeric heterochromatin via an HP1 interaction. J. Cell Sci. 115, 3439–3448 (2002).

5. Cammas, F., Herzog, M., Lerouge, T., Chambon, P. & Losson, R. Association of the transcriptional corepressor TIF1beta with heterochromatin protein 1 (HP1): an essential role for progression through differentiation. Genes Dev. 18, 2147–2160 (2004).

6. Friedman, J. R. et al. KAP-1, a novel corepressor for the highly conserved KRAB repression domain. Genes Dev. 10, 2067–2078 (1996).

7. Groner, A. C. et al. KRAB-zinc finger proteins and KAP1 can mediate long-range transcriptional repression through heterochromatin spreading. PLoS Genet. 6, e1000869 (2010).

8. Li, X. et al. Role for KAP1 serine 824 phosphorylation and sumoylation/desumoylation switch in regulating KAP1-mediated transcriptional repression. J. Biol. Chem. 282, 36177–36189 (2007).

9. Moosmann, P., Georgiev, O., Le Douarin, B., Bourquin, J. P. & Schaffner, W. Transcriptional repression by RING finger protein TIF1 beta that interacts with the KRAB repressor domain of KOX1. Nucleic Acids Res. 24, 4859–4867 (1996).

10. Bunch, H. et al. TRIM28 regulates RNA polymerase II promoter-proximal pausing and pause release. Nat. Struct. Mol. Biol. 21, 876–883 (2014).

11. Hu, G. et al. A genome-wide RNAi screen identifies a new transcriptional module required for selfrenewal. Genes Dev. 23, 837–848 (2009).

12. Iyengar, S., Ivanov, A. V., Jin, V. X., Rauscher, F. J. & Farnham, P. J. Functional Analysis of KAP1 Genomic Recruitment. Mol. Cell. Biol. 31, 1833–1847 (2011).

13. McNamara, R. P. et al. KAP1 Recruitment of the 7SK snRNP Complex to Promoters Enables Transcription Elongation by RNA Polymerase II. Mol. Cell 61, 39–53 (2016).

14. Goodarzi, A. A., Kurka, T. & Jeggo, P. A. KAP-1 phosphorylation regulates CHD3 nucleosome remodeling during the DNA double-strand break response. Nat. Struct. Mol. Biol. 18, 831–839 (2011).

15. Hu, C. et al. Roles of Kruppel-associated Box (KRAB)-associated Co-repressor KAP1 Ser-473 Phosphorylation in DNA Damage Response. J. Biol. Chem. 287, 18937–18952 (2012).

16. Li, X. et al. SUMOylation of the transcriptional co-repressor KAP1 is regulated by the serine and threonine phosphatase PP1. Sci. Signal. 3, ra32 (2010).

17. White, D. et al. The ATM substrate KAP1 controls DNA repair in heterochromatin: regulation by HP1 proteins and serine 473/824 phosphorylation. Mol. Cancer Res. MCR 10, 401–414 (2012).

18. White, D. E. et al. KAP1, a novel substrate for PIKK family members, colocalizes with numerous damage response factors at DNA lesions. Cancer Res. 66, 11594–11599 (2006).

19. Ziv, Y. et al. Chromatin relaxation in response to DNA double-strand breaks is modulated by a novel ATM- and KAP-1 dependent pathway. Nat. Cell Biol. 8, 870–876 (2006).

20. Bhatia, N. et al. Identification of novel small molecules that inhibit protein-protein interactions between MAGE and KAP-1. Arch. Biochem. Biophys. 508, 217–221 (2011).

21. Chen, L. et al. Tripartite Motif Containing 28 (Trim28) Can Regulate Cell Proliferation by Bridging HDAC1/E2F Interactions. J. Biol. Chem. 287, 40106–40118 (2012).

22. Jakobsson, J. et al. KAP1-mediated epigenetic repression in the forebrain modulates behavioral vulnerability to stress. Neuron 60, 818–831 (2008).

23. Lin, L.-F. et al. Loss of ZBRK1 contributes to the increase of KAP1 and promotes KAP1-mediated metastasis and invasion in cervical cancer. PloS One 8, e73033 (2013).

24. Okamoto, K., Kitabayashi, I. & Taya, Y. KAP1 dictates p53 response induced by chemotherapeutic agents via Mdm2 interaction. Biochem. Biophys. Res. Commun. 351, 216–222 (2006).

25. Wang, C. et al. MDM2 interaction with nuclear corepressor KAP1 contributes to p53 inactivation. EMBO J. 24, 3279–3290 (2005).

26. Yang, B. et al. MAGE-A, mMage-b, and MAGE-C proteins form complexes with KAP1 and suppress p53-dependent apoptosis in MAGE-positive cell lines. Cancer Res. 67, 9954–9962 (2007).

27. Yokoe, T. et al. KAP1 is associated with peritoneal carcinomatosis in gastric cancer. Ann. Surg. Oncol. 17, 821–828 (2010).

28. Ozato, K., Shin, D.-M., Chang, T.-H. & Morse, H. C. TRIM family proteins and their emerging roles in innate immunity. Nat. Rev. Immunol. 8, 849–860 (2008).

29. Saurin, A. J., Borden, K. L., Boddy, M. N. & Freemont, P. S. Does this have a familiar RING? Trends Biochem. Sci. 21, 208–214 (1996).

30. Doyle, J. M., Gao, J., Wang, J., Yang, M. & Potts, P. R. MAGE-RING protein complexes comprise a family of E3 ubiquitin ligases. Mol. Cell 39, 963–974 (2010).

31. Liang, Q. et al. Tripartite motif-containing protein 28 is a small ubiquitin-related modifier E3 ligase and negative regulator of IFN regulatory factor 7. J. Immunol. Baltim. Md 1950 187, 4754–4763 (2011).

32. Rousseaux, M. W. et al. Depleting Trim28 in adult mice is well tolerated and reduces levels of α-synuclein and tau. eLife 7, (2018).

33. Meroni, G. & Diez-Roux, G. TRIM/RBCC, a novel class of ‘single protein RING finger’ E3 ubiquitin ligases. BioEssays News Rev. Mol. Cell. Dev. Biol. 27, 1147–1157 (2005).

34. Peng, H. et al. Reconstitution of the KRAB-KAP-1 repressor complex: a model system for defining the molecular anatomy of RING-B box-coiled-coil domain-mediated protein-protein interactions. J. Mol. Biol. 295, 1139–1162 (2000).

35. Ragvin, A. et al. Nucleosome binding by the bromodomain and PHD finger of the transcriptional cofactor p300. J. Mol. Biol. 337, 773–788 (2004).

36. Zhou, Y. & Grummt, I. The PHD finger/bromodomain of NoRC interacts with acetylated histone H4K16 and is sufficient for rDNA silencing. Curr. Biol. CB 15, 1434–1438 (2005).

37. Ivanov, A. V. et al. PHD domain-mediated E3 ligase activity directs intramolecular sumoylation of an adjacent bromodomain required for gene silencing. Mol. Cell 28, 823–837 (2007).

38. Neo, S. H. et al. TRIM28 Is an E3 Ligase for ARF-Mediated NPM1/B23 SUMOylation That Represses Centrosome Amplification. Mol. Cell. Biol. 35, 2851–2863 (2015).

39. Yang, Y. et al. Acetylated hsp70 and KAP1-mediated Vps34 SUMOylation is required for autophagosome creation in autophagy. Proc. Natl. Acad. Sci. U. S. A. 110, 6841–6846 (2013).

40. Zeng, L. et al. Structural insights into human KAP1 PHD finger-bromodomain and its role in gene silencing. Nat. Struct. Mol. Biol. 15, 626–633 (2008).

41. Ryan, R. F. et al. KAP-1 corepressor protein interacts and colocalizes with heterochromatic and euchromatic HP1 proteins: a potential role for Krüppel-associated box-zinc finger proteins in heterochromatin-mediated gene silencing. Mol. Cell. Biol. 19, 4366–4378 (1999).

42. Schultz, D. C., Friedman, J. R. & Rauscher, F. J. Targeting histone deacetylase complexes via KRAB-zinc finger proteins: the PHD and bromodomains of KAP-1 form a cooperative unit that recruits a novel isoform of the Mi-2α subunit of NuRD. Genes Dev. 15, 428–443 (2001).

43. Schultz, D. C., Ayyanathan, K., Negorev, D., Maul, G. G. & Rauscher, F. J. SETDB1: a novel KAP-1-associated histone H3, lysine 9-specific methyltransferase that contributes to HP1-mediated silencing of euchromatic genes by KRAB zinc-finger proteins. Genes Dev. 16, 919–932 (2002).

44. Sripathy, S. P., Stevens, J. & Schultz, D. C. The KAP1 corepressor functions to coordinate the assembly of de novo HP1-demarcated microenvironments of heterochromatin required for KRAB zinc finger protein-mediated transcriptional repression. Mol. Cell. Biol. 26, 8623–8638 (2006).

45. Underhill, C., Qutob, M. S., Yee, S. P. & Torchia, J. A novel nuclear receptor corepressor complex, N-CoR, contains components of the mammalian SWI/SNF complex and the corepressor KAP-1. J. Biol. Chem. 275, 40463–40470 (2000).

46. Lechner, M. S., Begg, G. E., Speicher, D. W. & Rauscher, F. J. Molecular Determinants for Targeting Heterochromatin Protein 1-Mediated Gene Silencing: Direct Chromoshadow Domain–KAP-1 Corepressor Interaction Is Essential. Mol. Cell. Biol. 20, 6449–6465 (2000).

47. Smothers, J. F. & Henikoff, S. The HP1 chromo shadow domain binds a consensus peptide pentamer. Curr. Biol. CB 10, 27–30 (2000).

48. Eissenberg, J. C. & Elgin, S. C. R. HP1a: a structural chromosomal protein regulating transcription. Trends Genet. TIG 30, 103–110 (2014).

49. Nishibuchi, G. & Nakayama, J. Biochemical and structural properties of heterochromatin protein 1: understanding its role in chromatin assembly. J. Biochem. (Tokyo) 156, 11–20 (2014).

50. Thiru, A. et al. Structural basis of HP1/PXVXL motif peptide interactions and HP1 localisation to heterochromatin. EMBO J. 23, 489–499 (2004).

51. Chang, C.-W. et al. Phosphorylation at Ser473 regulates heterochromatin protein 1 binding and corepressor function of TIF1 beta/KAP1. BMC Mol. Biol. 9, 61 (2008).

52. Huntley, S. et al. A comprehensive catalog of human KRAB-associated zinc finger genes: insights into the evolutionary history of a large family of transcriptional repressors. Genome Res. 16, 669–677 (2006).

53. Imbeault, M., Helleboid, P.-Y. & Trono, D. KRAB zinc-finger proteins contribute to the evolution of gene regulatory networks. Nature 543, 550–554 (2017).

54. Urrutia, R. KRAB-containing zinc-finger repressor proteins. Genome Biol. 4, 231 (2003).

55. Koliopoulos, M. G. et al. Molecular mechanism of influenza A NS1-mediated TRIM25 recognition and inhibition. Nat. Commun. 9, 1820 (2018).

56. Sanchez, J. G. et al. The tripartite motif coiled-coil is an elongated antiparallel hairpin dimer. Proc. Natl. Acad. Sci. U. S. A. 111, 2494–2499 (2014).

57. Goldstone, D. C. et al. Structural studies of postentry restriction factors reveal antiparallel dimers that enable avid binding to the HIV-1 capsid lattice. Proc. Natl. Acad. Sci. U. S. A. 111, 9609–9614 (2014).

58. Wagner, J. M. et al. Mechanism of B-box 2 domain-mediated higher-order assembly of the retroviral restriction factor TRIM5α. eLife 5, (2016).

59. Weinert, C., Morger, D., Djekic, A., Grütter, M. G. & Mittl, P. R. E. Crystal structure of TRIM20 C-terminal coiled-coil/B30.2 fragment: implications for the recognition of higher order oligomers. Sci. Rep. 5, 10819 (2015).

60. Li, Y. et al. Structural insights into the TRIM family of ubiquitin E3 ligases. Cell Res. 24, 762–765 (2014).

61. Koliopoulos, M. G., Esposito, D., Christodoulou, E., Taylor, I. A. & Rittinger, K. Functional role of TRIM E3 ligase oligomerization and regulation of catalytic activity. EMBO J. 35, 1204–1218 (2016).

62. Herquel, B. et al. Transcription cofactors TRIM24, TRIM28, and TRIM33 associate to form regulatory complexes that suppress murine hepatocellular carcinoma. Proc. Natl. Acad. Sci. U. S. A. 108, 8212–8217 (2011).

63. Li, X., Yeung, D. F., Fiegen, A. M. & Sodroski, J. Determinants of the higher order association of the restriction factor TRIM5alpha and other tripartite motif (TRIM) proteins. J. Biol. Chem. 286, 27959–27970 (2011).

64. Svergun, D. I., Petoukhov, M. V. & Koch, M. H. Determination of domain structure of proteins from X-ray solution scattering. Biophys. J. 80, 2946–2953 (2001).

65. Grudinin, S., Garkavenko, M. & Kazennov, A. Pepsi-SAXS: an adaptive method for rapid and accurate computation of small-angle X-ray scattering profiles. Acta Crystallogr. Sect. Struct. Biol. 73, 449–464 (2017).

66. Hoffmann, A. & Grudinin, S. NOLB: Nonlinear Rigid Block Normal-Mode Analysis Method. J. Chem. Theory Comput. 13, 2123–2134 (2017).

67. Ortega, A., Amorós, D. & García de la Torre, J. Prediction of hydrodynamic and other solution properties of rigid proteins from atomic- and residue-level models. Biophys. J. 101, 892–898 (2011).

68. Svergun, D. I. Restoring Low Resolution Structure of Biological Macromolecules from Solution Scattering Using Simulated Annealing. Biophys. J. 76, 2879–2886 (1999).

69. Grant, T. D. Ab initio electron density determination directly from solution scattering data. Nat. Methods 15, 191–193 (2018).

70. Degiacomi, M. T. et al. Molecular assembly of the aerolysin pore reveals a swirling membrane-insertion mechanism. Nat. Chem. Biol. 9, 623–629 (2013).

71. Tamò, G. et al. Disentangling constraints using viability evolution principles in integrative modeling of macromolecular assemblies. Sci. Rep. 7, 235 (2017).

72. Tamò, G. E., Abriata, L. A. & Dal Peraro, M. The importance of dynamics in integrative modeling of supramolecular assemblies. Curr. Opin. Struct. Biol. 31, 28–34 (2015).

73. Bernardi, R. & Pandolfi, P. P. Structure, dynamics and functions of promyelocytic leukaemia nuclear bodies. Nat. Rev. Mol. Cell Biol. 8, 1006–1016 (2007).

74. Shen, T. H., Lin, H.-K., Scaglioni, P. P., Yung, T. M. & Pandolfi, P. P. The mechanisms of PML-nuclear body formation. Mol. Cell 24, 331–339 (2006).

75. Schindelin, J. et al. Fiji: an open-source platform for biological-image analysis. Nat. Methods 9, 676–682 (2012).

76. Bryan, L. C. et al. Single-molecule kinetic analysis of HP1-chromatin binding reveals a dynamic network of histone modification and DNA interactions. Nucleic Acids Res. 45, 10504–10517 (2017).

77. Kilic, S., Bachmann, A. L., Bryan, L. C. & Fierz, B. Multivalency governs HP1α association dynamics with the silent chromatin state. Nat. Commun. 6, 7313 (2015).

78. Chi, P., Allis, C. D. & Wang, G. G. Covalent histone modifications--miswritten, misinterpreted and mis-erased in human cancers. Nat. Rev. Cancer 10, 457–469 (2010).

79. Ruthenburg, A. J., Li, H., Patel, D. J. & Allis, C. D. Multivalent engagement of chromatin modifications by linked binding modules. Nat. Rev. Mol. Cell Biol. 8, 983–994 (2007).

80. Smith, E. & Shilatifard, A. The chromatin signaling pathway: diverse mechanisms of recruitment of histone-modifying enzymes and varied biological outcomes. Mol. Cell 40, 689–701 (2010).

81. Matsuoka, S. et al. ATM and ATR substrate analysis reveals extensive protein networks responsive to DNA damage. Science 316, 1160–1166 (2007).

82. D’Cruz, A. A. et al. Identification of a second binding site on the TRIM25 B30.2 domain. Biochem. J. 475, 429–440 (2018).

83. Dawidziak, D. M., Sanchez, J. G., Wagner, J. M., Ganser-Pornillos, B. K. & Pornillos, O. Structure and catalytic activation of the TRIM23 RING E3 ubiquitin ligase. Proteins 85, 1957–1961 (2017).

84. Larson, A. G. et al. Liquid droplet formation by HP1α suggests a role for phase separation in heterochromatin. Nature 547, 236–240 (2017).

85. Hendriks, I. A. et al. Site-specific mapping of the human SUMO proteome reveals co-modification with phosphorylation. Nat. Struct. Mol. Biol. 24, 325–336 (2017).

86. Marcaida, M. J., DePristo, M. A., Chandran, V., Carpousis, A. J. & Luisi, B. F. The RNA degradosome: life in the fast lane of adaptive molecular evolution. Trends Biochem. Sci. 31, 359–365 (2006).

87. Schuck, P. Size-distribution analysis of macromolecules by sedimentation velocity ultracentrifugation and lamm equation modeling. Biophys. J. 78, 1606–1619 (2000).

88. Laue, T., Shaw, B. D., Ridgeway, T. M., and Pelletier, S. L. Analytical Ultracentrifugation in Biochemistry and Polymer Science. in 90–125 (The Royal Society of Chemistry, Cambridge, U.K., 1992).

89. Incardona, M.-F. et al. EDNA: a framework for plugin-based applications applied to X-ray experiment online data analysis. J. Synchrotron Radiat. 16, 872–879 (2009).

90. Franke, D. et al. ATSAS 2.8: a comprehensive data analysis suite for small-angle scattering from macromolecular solutions. J. Appl. Crystallogr. 50, 1212–1225 (2017).

91. Rambo, R. P. Resolving Individual Components in Protein-RNA Complexes Using Small-Angle X-ray Scattering Experiments. Methods Enzymol. 558, 363–390 (2015).

92. Panjkovich, A. & Svergun, D. I. CHROMIXS: automatic and interactive analysis of chromatography-coupled small-angle X-ray scattering data. Bioinforma. Oxf. Engl. 34, 1944–1946 (2018).

93. Konarev, P. V., Volkov, V. V., Sokolova, A. V., Koch, M. H. J. & Svergun, D. I. PRIMUS: a Windows PC-based system for small-angle scattering data analysis. J. Appl. Crystallogr. 36, 1277–1282 (2003).

94. Svergun, D. I. Determination of the regularization parameter in indirect-transform methods using perceptual criteria. J. Appl. Crystallogr. 25, 495–503 (1992).

95. Humphrey, W., Dalke, A. & Schulten, K. VMD: visual molecular dynamics. J. Mol. Graph. 14, 33–38, 27–28 (1996).

96. Pettersen, E. F. et al. UCSF Chimera--a visualization system for exploratory research and analysis. J. Comput. Chem. 25, 1605–1612 (2004).

97. Biasini, M. et al. SWISS-MODEL: modelling protein tertiary and quaternary structure using evolutionary information. Nucleic Acids Res. 42, W252–258 (2014).

98. Huang, A. et al. Symmetry and Asymmetry of the RING–RING Dimer of Rad18. J. Mol. Biol. 410, 424–435 (2011).

99. Mrosek, M. et al. Structural Analysis of B-Box 2 from MuRF1: Identification of a Novel Self-Association Pattern in a RING-like Fold. Biochemistry 47, 10722–10730 (2008).

100. Kim, D. E., Chivian, D. & Baker, D. Protein structure prediction and analysis using the Robetta server. Nucleic Acids Res. 32, W526–W531 (2004).

101. Sali, A. & Blundell, T. L. Comparative protein modelling by satisfaction of spatial restraints. J. Mol. Biol. 234, 779–815 (1993).

102. Huang, Y., Myers, M. P. & Xu, R.-M. Crystal structure of the HP1-EMSY complex reveals an unusual mode of HP1 binding. Struct. Lond. Engl. 1993 14, 703–712 (2006).

103. Greber, B. J. et al. The complete structure of the large subunit of the mammalian mitochondrial ribosome. Nature 515, 283–286 (2014).

104. Leaver-Fay, A. et al. Rosetta3: An Object-Oriented Software Suite for the Simulation and Design of Macromolecules. Methods Enzymol. 487, 545–574 (2011).

105. Scheres, S. H. W. RELION: implementation of a Bayesian approach to cryo-EM structure determination. J. Struct. Biol. 180, 519–530 (2012).

106. Vila-Perelló, M. et al. Streamlined expressed protein ligation using split inteins. J. Am. Chem. Soc. 135, 286–292 (2013).

## References

1. Schuck, P. Size-distribution analysis of macromolecules by sedimentation velocity ultracentrifugation and lamm equation modeling. Biophys. J. 78, 1606–1619 (2000).

2. Konarev, P. V., Volkov, V. V., Sokolova, A. V., Koch, M. H. J. & Svergun, D. I. PRIMUS: a Windows PC-based system for small-angle scattering data analysis. J Appl Cryst, J Appl Crystallogr 36, 1277–1282 (2003).

3. Rambo, R. P. Resolving Individual Components in Protein-RNA Complexes Using Small-Angle X-ray Scattering Experiments. Meth. Enzymol. 558, 363–390 (2015).

4. Bernadó, P., Mylonas, E., Petoukhov, M. V., Blackledge, M. & Svergun, D. I. Structural Characterization of Flexible Proteins Using Small-Angle X-ray Scattering. Journal of the American Chemical Society 129, 5656–5664 (2007).

